# The Phage-shock-protein (PSP) Envelope Stress Response: Discovery of Novel Partners and Evolutionary History

**DOI:** 10.1101/2020.09.24.301986

**Authors:** Janani Ravi, Vivek Anantharaman, Samuel Zorn Chen, Evan Pierce Brenner, Pratik Datta, L Aravind, Maria Laura Gennaro

## Abstract

Bacterial phage shock protein (PSP) systems stabilize the bacterial cell membrane and protect against envelope stress. These systems have been associated with virulence, but despite their critical roles, PSP components are not well-characterized outside proteobacteria. Using comparative genomics and protein sequence-structure-function analyses, we systematically identified and analyzed PSP homologs, phyletic patterns, domain architectures, and gene neighborhoods. This approach underscored the evolutionary significance of the system, revealing that the core PspA gene (Snf7 in ESCRT outside bacteria) was present in the Last Universal Common Ancestor (LUCA), and that this ancestral functionality has since diversified into multiple novel, distinct PSP systems across life. Several novel partners of the PSP system were identified: (i) the Toastrack domain, likely facilitating assembly of sub-membrane stress-sensing and signaling complexes, (ii) the newly-defined HAAS-PadR-like transcriptional regulator pair system, and (iii) multiple independent associations with ATPase, CesT/Tir-like chaperone, and Band-7 domains in proteins thought to mediate sub-membrane dynamics. Our work also uncovered links between the PSP components and other domains, such as novel variants of SHOCT-like domains, suggesting roles in assembling membrane-associated complexes of proteins with disparate biochemical functions. Results are available at https://jravilab.org/psp.

**Importance:** Phage shock proteins (PSP) are virulence-associated, cell membrane stress-protective systems. They have mostly been characterized in proteobacteria and firmicutes. We now show that PSP systems were present in the Last Universal Common Ancestor, and that homologs have evolved and diversified into newly identified functional contexts. Recognizing the conservation and evolution of PSP systems across bacterial phyla contributes to our understanding of stress response mechanisms in prokaryotes. Moreover, the newly discovered PSP modularity will likely prompt new studies of lineage-specific cell-envelope structures, lifestyles, and adaptation mechanisms. Finally, our results validate use of domain architecture and genetic context for discovery in comparative genomics.

## Introduction

Cell membranes are complex, dynamic structures made of bilayer and non-bilayer lipids and proteins (1). These membrane components engage in critical processes, including membrane biogenesis, cell shape maintenance and division, small molecule transport, inter- and intracellular signaling, maintenance of proton motive force (PMF), and motility through cytoskeletal proteins (2). Cell membranes continuously adapt to external stresses, particularly in unicellular organisms (3–7). Failure to adjust leads to membrane damage and cell death. A specialized suite of lipids and proteins maintain membrane form and function under stress (8, 9), including the multiprotein bacterial Phage Shock Protein (PSP) system, and the archaeo-eukaryotic ESCRT system (10–20).

The origins and divergence of membrane structure and function remain open questions in the very basis of life. Most membrane functions are conserved across the tree of life despite compositional and structural variations (1), which provides insights into evolutionary history and niche specialization. Where functions differ can also be informative. Prior research has validated that evolutionary processes across life are constrained, and the potential evolutionary paths are not limitless (21, 22). Mapping out evolutionary history is, therefore, more attainable by the existence of constrained characteristics – a basis for phenotypic convergence. Membrane maintenance mechanisms obey these same rules, and characterizing membrane homeostasis systems like PSP can provide functional and evolutionary insights across life.

PSP functionality is driven by the peripheral membrane effector protein PspA, and membrane stability phenotypes are restored in *E. coli* Δ*pspA* strains by complementation with the homologous plant protein, Vipp1 (11–16, 23–31). Both proteins contain the coiled-coil structure PspA domain, and both have been shown to form large homooligomeric complexes (16, 17, 19, 20, 32, 33) (**Table S1**). Bacterial PspA senses and responds to extracellular stress (14, 23, 33–35). Its domain, Pfam PspA_IM30 (inner membrane 30KDa domain) is found in bacteria, photosynthetic eukaryotes, and archaea (15–17, 33, 36, 37) (**Fig. 1A**). Proteins containing PspA_IM30 are known to facilitate membrane fusion (16) and prevent leakage of protons through the membrane (2, 23) in response to a wide range of surface stresses, including changes in membrane potential, *ΔpH* (2, 23) and PMF (38–41), (l)antibiotic stress (7), heat/osmotic shock (42) and mechanical stress via stored curvature elastic stress (13, 43–45). Vipp1 is involved in thylakoid biogenesis, vesicular lipid transport and storage, and membrane shaping (16, 26, 27, 32).

**Figure 1.**
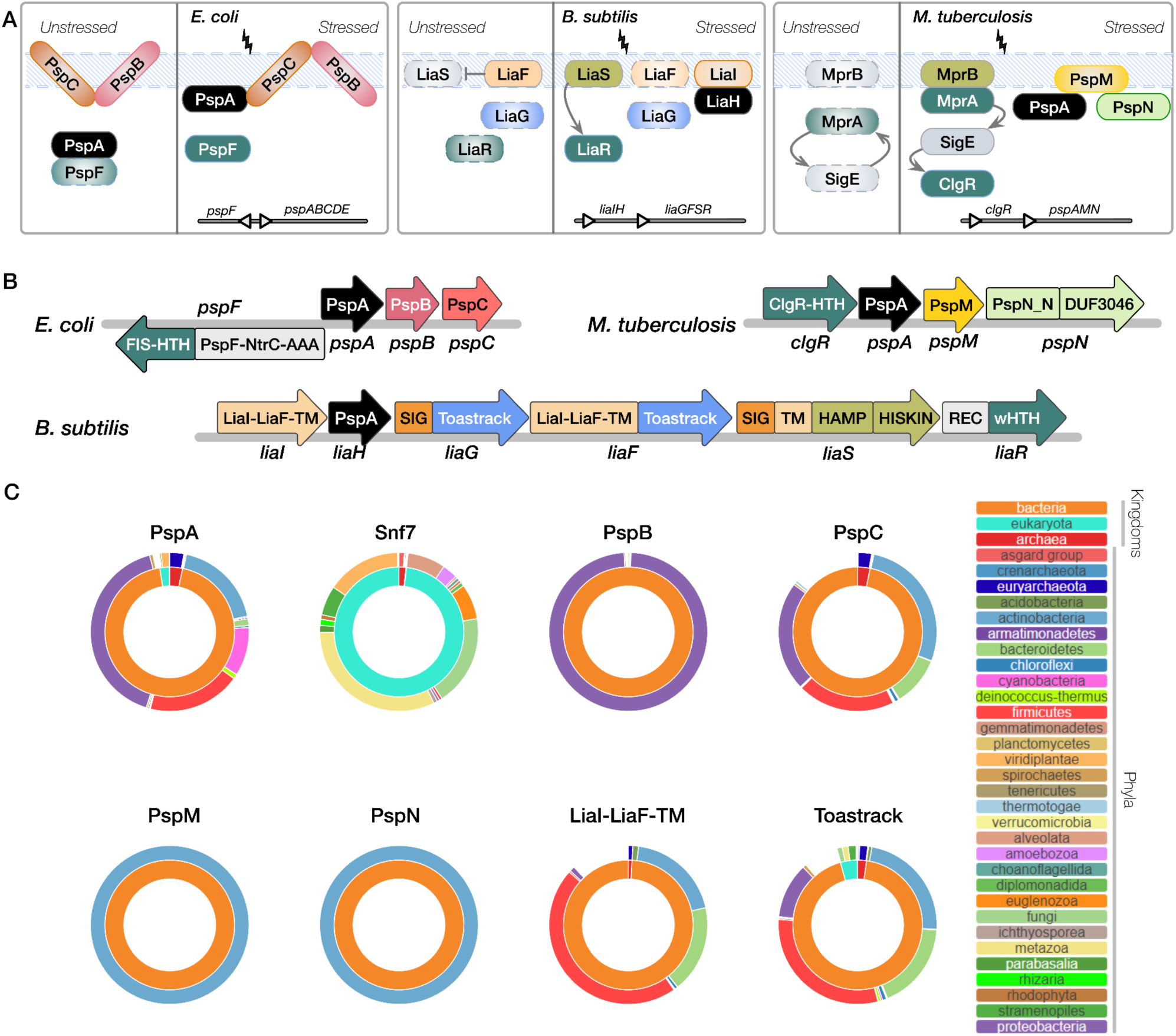
The phyletic spread of classical PSP members across all major lineages. **A. The three classical PSP systems** in *E. coli* (*pspF*||*pspABC*), *M. tuberculosis* (*clgRpspAMN*), and **B.** *subtilis* (*liaIHGFSR*) are shown. Proteins across three bacterial PSP systems are labeled and colors uniformly (across all figures that show domain architectures) indicate the nature of the protein; ***black***, PspA homolog (*e.g.,* PspA, LiaH); ***teal***, transcription factor/response regulator (*e.g.,* PspF, LiaR, MprA, ClgR) along with partner histidine kinases in ***olive green*** (*e.g.,* LiaS, MprB part of two-component systems LiaRS, MprAB); ***orange/yellow***, transmembrane protein (*e.g.,* PspB, PspC, LiaF, LiaI, PspM) embedded in the dashed blue membrane. Arrows represent interaction or activation, or inhibitory feedback in the case of LiaF and LiaS. A simple operon map is shown for each system at the bottom, but is expanded upon in panel **B**. **B. Domain architectures of the classical PSP operons** in *E. coli*, *M. tuberculosis*, and *B. subtilis*. Domains are denoted by rectangular segments inside the block arrow representing each protein labeled below the arrow (*e.g.*, the protein LiaH contains only the PspA domain). The direction of the arrow indicates the direction of transcription. See *Results* for new domain definitions. **C. Phyletic spreads of PSP proteins.** Sunburst plots are shown to indicate the lineage distributions (as a fraction of the total number of homologs recorded across lineages) for the homologs of ‘classical’ domains/protein families of interest: PspA, Snf7, PspB, PspC, PspM, PspN (and DUF3046), LiaI-LiaF-TM, Toastrack. In each plot corresponding to a particular protein, the inner ring corresponds to the proportion of its homologs present in superkingdoms/domains of life, Bacteria, Archaea, and Eukaryota. The outer ring depicts the distribution of the homologs among key phyla. Interactive sunburst plots for each of the PSP proteins are available in the <u>web app</u>. The colors for each lineage (outer and inner rings) are shown separately for **C**.

The classical PSP system first discovered in Proteobacteria (46) is not conserved across all bacterial and archaeal clades (12, 33, 46). However, the PSP systems described in Firmicutes and Actinobacteria (3, 11, 14, 15, 35, 36, 47, 48) include clade-specific PSP proteins with distinct domain architectures and conserved functions (*e.g.,* transcriptional regulation, membrane tethering and relocation). Several key questions remain, including how PspA evolved to function with distinct partners across diverse lineages, about the identities of other integral components of the PSP system beyond the classical systems and complete characterization of cognate partners, if any homologs participate in novel functions, and whether common themes emerge from the phyletic patterns of PSP systems.

We set out to evaluate the evolutionary and functional significance of the PSP system through a comprehensive analysis of PspA occurrences, associated genomic contexts, and phyletic spreads across the three superkingdoms or domains of life (49, 50). To perform this, we resolved PSP proteins into their constituent domain architectures and queried each individual domain in our comparative genomics and phylogenetic analyses. This domain-centric approach identified several novel players in the PSP envelope stress response system, of which we described putative function and evolution. We have also made available the entire PSP repertoire — including new members, affiliate proteins, homologs (including paralogs and orthologs), domain architectures, genomic neighborhoods, phyletic spreads, and phylogenetic analyses in all representative species — for researchers through tabular and graphical summaries on our user-friendly, interactive companion <u>web application</u>.

## Results and Discussion

PspA is found widely across the tree of life in distinct genomic neighborhoods and with different functional partners (15, 30, 51). Using comparative genomics and a computational molecular evolutionary approach, we therefore sought to comprehensively identify and characterize all homologs of PspA and cognate partner domains across a dataset representative of the tree of life. We discovered several novel conserved, lineage-specific domain architectures and genomic contexts of PSP systems and delineated their phyletic spreads across the tree of life in the process (see *Methods* for details). All our findings (data, results, and visual summaries) are available in an interactive and searchable web application (https://jravilab.org/psp) that can be queried with the protein accession numbers cited inline throughout the *Results* section.

### Phyletic spread of classical PSP components across the tree of life

To describe the network of PSP proteins across the tree of life we analyzed the classical PSP components in Proteobacteria, Actinobacteria, and Firmicutes. Among Proteobacteria, we traced the *E. coli psp* operon, which is widely conserved across Gammaproteobacteria (10, 12, 39, 52–55). PspA, PspB, and PspC are each composed of their namesake domains spanning almost their entire protein lengths. The transcriptional regulator PspF is encoded on the opposite strand and shares a central promoter region with the *pspABC* operon in most Gammaproteobacteria, and only with *pspA* in other organisms. PspF is an enhancer-binding protein, with a NtrC-type AAA+ domain fused to a C-terminal Fis-type HTH (helix-turn-helix) DNA-binding domain. In Actinobacteria, the *M. tuberculosis psp* operon (14, 15, 36) includes the namesake PspA protein; a transcription factor, ClgR, unique to Actinobacteria and containing a Cro-like-HTH (cHTH) DNA-binding domain; an integral membrane protein, Rv2743c, distinct from *E. coli* PspC and PspB; and Rv2742c, an uncharacterized protein carrying a “Domain of Unknown Function,’’ <u>DUF3046</u> (14, 15, 36). For consistency, we rename Rv2743c and Rv2742c as PspM and PspN, respectively (15). *B. subtilis,* a member of Firmicutes, contains the *lia* operon. LiaI is a small transmembrane (TM) protein (11, 48, 56), LiaF contains N-terminal TM helices (<u>DUF2157</u>) and a C-terminal globular domain (<u>DUF2154</u>), and LiaG contains an uncharacterized domain <u>DUF4097</u>. We defined two novel domains of the PSP system in these operons: the LiaI-LiaF-TM sensory domain that unifies the four TM domains (4TM) and <u>DUF2157</u> from LiaI and LiaF, and Toastrack, unifying <u>DUF4097</u>/<u>DUF2154</u>/<u>DUF2807</u> Pfam models. The *lia* operon is controlled by a two-component system LiaRS (11, 35). LiaR contains a receiver domain, and a winged HTH (wHTH) DNA-binding domain (REC-wHTH), while LiaS is a sensor kinase with a TM region, intracellular HAMP, and a histidine kinase (HISKIN) signaling domain (57). Information on domains in this section is presented in **Text S1.1** and **Table S1**, and graphically in **Fig. 1A** and **1B**.

### PspA

From six distinct queries (Proteobacteria, Firmicutes, Actinobacteria, Cyanobacteria, Plants; see *Methods*), homologs of PspA were sought comprehensively. Consistent with prior studies, PspA_IM30 was identified in most bacterial and archaeal lineages (**Fig. 1C**; <u>web app</u>). Within eukaryotes, only the SAR and Archaeplastida (greater plant lineage) carry PspA_Vipp1 homologs (**Fig. 1C**). Analysis using multi-query sequence searches, we also identified Snf7 proteins from the eukaryotic membrane-remodeling ESCRT systems (14, 18, 58–61) as homologs (**Fig. 1B**). Snf7 was then used for iterative searches, which yielded several pan-eukaryotic and archaeal homologs distinct from Vipp1 (**Fig. 1B** and **1C**; **Table S3**), supporting a considerably wider evolutionary distribution for PspA, corroborated by other recent reports (19, 20).

### Toastrack and LiaI-LiaF-TM

Iterative domain-centric searches and multiple sequence alignments (MSAs) revealed that LiaI and LiaG were similar to N-and C-terminal regions of LiaF. We define these two novel domains as **Toastrack** (unifying DUF2154, DUF4097, DUF2807) and **LiaI-LiaF-TM**. Toastrack-containing proteins are characterized by a single-stranded right-handed beta-helix fold with a unique N-terminal 7-stranded region featuring complex intertwining of the beta-helix strands (e.g., PDB: <u>4QRK</u>, <u>4OPW</u>). They are pan-bacterial and sporadic in archaea and eukaryotes (**Fig. 1C**). Toastrack is frequently coupled to PspC, LiaI-LiaF-TM, or both (**Fig. 4**; **Table S4**). LiaI-LiaF-TM contains a 4TM region, and is primarily in bacteria, with transfers to Euryarchaeota (**Fig. 1B**). Search methodology is described in **Text S1.1**.

### PspB and PspC

**PspB**, often part of the proteobacterial *psp* operon, is another integral membrane protein implicated in stress sensing and virulence (62–64). PspB has an N-terminal anchoring TM helix followed by an α-helical cytoplasmic globular domain. While PspB is rarely found outside proteobacteria (**Fig. 1C**), we discovered previously unrecognized divergent PspB domains fused to PspC (see section on *PspC architectures)*.

**PspC** was first identified in the proteobacterial PSP system; it is critical in sensing membrane stress and restoring envelope integrity. Recent studies demonstrate that PspC in some bacterial species may function independently of PspA in response to certain membrane stressors (58, 63, 65, 66). Another integral membrane protein, our analysis showed PspC has two TM helices, the first being a cryptic TM. We observed that the PspC domain is pan-bacterial and also present in a few archaeal clades (**Fig. 1C**). Our analysis showed that PspC has a conserved arginine between two TM helices, the first being a cryptic TM. We also identified several novel domain architectures and genomic contexts of PspC (see section on *PspC architectures*).

### PspM and PspN

Domain-centric searches found no discernible homologs for PspMN outside Actinobacteria (**Fig. 1C**; <u>web app</u>), as with prior full protein searches (15, 36, 67). Two TM helices make up the corynebacterial integral membrane partner **PspM** (Rv2743c). Our analyses revealed minimal PspM variation by MSA and a narrow phyletic spread restricted to mycobacteria and corynebacteria (**Fig. 1B**) (15, 36, 67).

The fourth member in the Mycobacterial operon, PspN (Rv2742c), contains a C-terminal <u>DUF3046</u> and an N-terminal domain we call **PspN_N** (**Table S1**; **Text S1.2**) (15, 36, 67). **DUF3046** is prevalent only in Actinobacteria (**Fig. 1C**). Moreover, the *M. tuberculosis* genome carries a second copy of DUF3046 (Rv2738c), located four genes downstream of *pspN*. We infer that Rv2738c, rather than *pspN*, contains the ancestral copy of DUF3046 (**Text S1.2**). DUF3046 found as part of PspN is likely a duplicate translocated into the PspN open reading frame (ORF) of mycobacteria, especially the *M. tuberculosis* complex. Further, unlike the pan-actinobacterial DUF3046, the N-terminal region, PspN_N, is conserved only in *M. tuberculosis*, with remnants of the coding region existing as potential pseudo-genes or degraded sequences in a few closely related mycobacteria (*e.g., M. avium*).

### Evolution of PspA

Since our results support recent findings that eukaryotic Snf7 is a homolog of PspA (19, 20), we used a structure-informed MSA and phylogenetic tree (‘Phylogeny’ tab, <u>web app</u>; **Fig. 2; Table S1**) to investigate the PspA/Snf7 superfamily and define its evolutionary origins.

**Figure 2.**
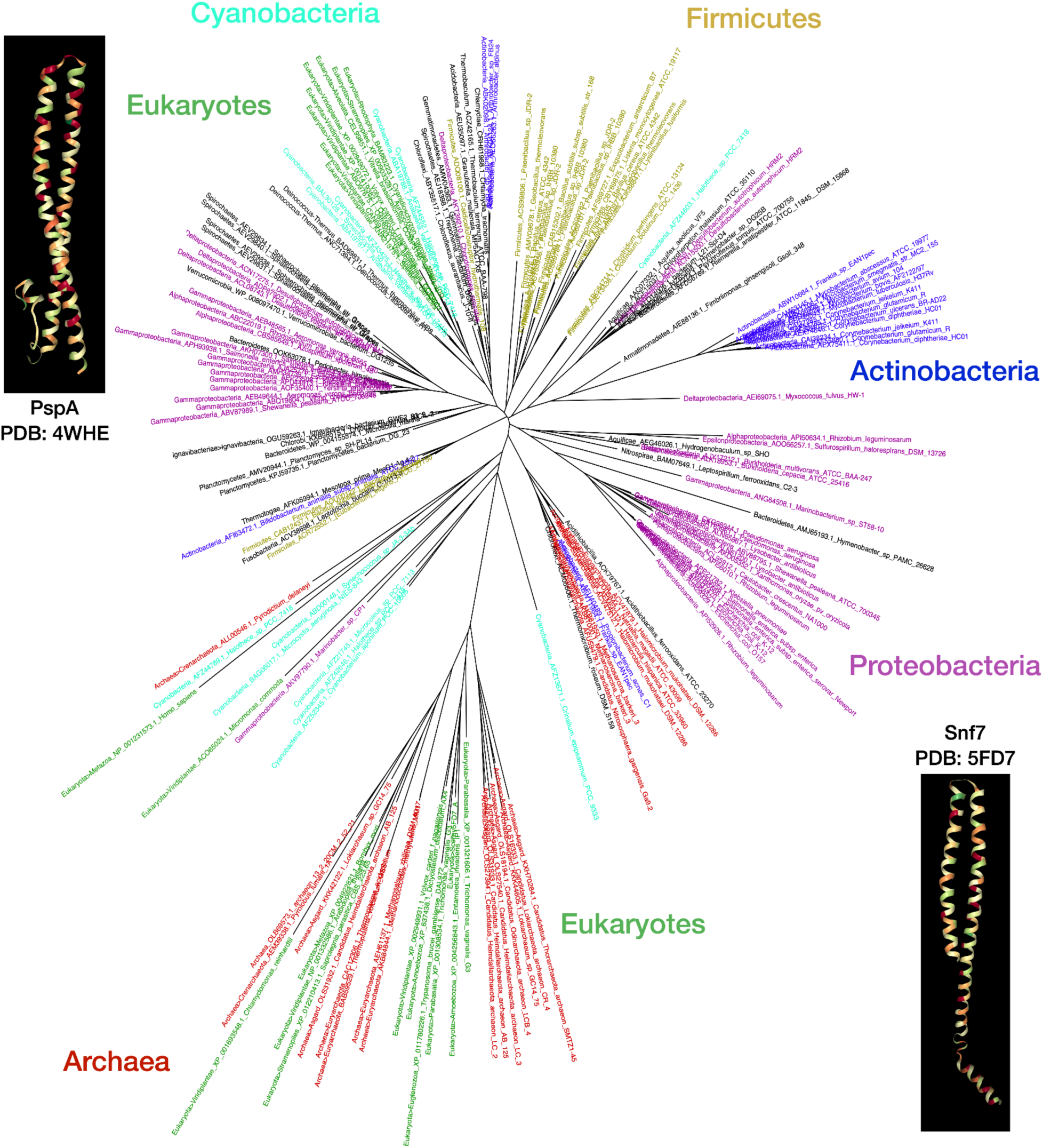
PspA/Snf7 homologs across the tree of life. **Phylogenetic tree of PspA homologs across the tree of life.** The phylogenetic tree was constructed using FastTree and visualized with FigTree (parallel RAxML-NG generated PspA tree with confidence values is shown in **Fig. S2**) from a multiple sequence alignment (using kalign3) of representative PspA homologs across all major superkingdoms and phyla (see *Methods*; <u>web app</u>). The primary takehome from the tree is not the 100s of leaves/labels but the key lineages that are labeled next to distinct clusters of similar PspA proteins; leaf colors match the lineage labels to highlight this import. The insets show the 3D structures for PspA (<u>4WHE</u>) and Snf7 (<u>5FD7</u>) from the Protein Data Bank. The tree leaves are labeled by lineage, species, and accession numbers. A text-searchable, scaled vector graphics version of this tree is available through the <u>web app</u> under the Phylogeny tab (leaf labels with aforementioned lineage, species, accession can query this high-resolution downloadable PDF/SVG version of the tree). Representatives for PspA/Snf7 homologs are also available in **Table S3**. As a gene tree, most homologs do cluster among evolutionarily similar species, though evidence of widespread horizontal gene transfer is also evident. Two clusters of eukaryotic genes appear: Snf7 in Eukaryota and Asgardarchaeota groups, mirroring recent research suggesting eukaryotic life arose from this clade; and Vipp1 among Eukaryota and Cyanobacteria group, again reflecting the shared origin of the protein in plants from Cyanobacteria.

#### Identifying clades carrying PspA and Snf7

In our analyses, all versions of the PspA domain phylogenetically more similar to the ‘classical’ PspA domain than the Snf7 domains were termed PspA+. We found that most bacterial lineages contain PspA+ clade members (**Fig. 1C**), with a few transfers to archaea. Among eukaryotes, only those containing plastids and two flagella (comprising the SAR/HA supergroup and excavata) have members of this clade (**Fig. 1C**; **Fig. 2**; <u>web app</u>; PDF version of the tree is searchable by the unique leaf identifiers carrying the accession, species, phylum) (59, 60).

We found that the curated set of PspA-like proteins contains a divergent cluster of Snf7 proteins in the ESCRT-III family (Snf7 family on Pfam/Interpro, part of PspA clan; **Table S1**), which we refer to as the Snf7+ clade (**Fig 2**; <u>web app</u>). The ESCRT-III complex is required for endosome-mediated trafficking via multivesicular body formation and sorting and has a predominantly archaeo-eukaryotic phyletic pattern (**Fig. 1C**) (18, 61, 68–71).

#### Tracing the roots of the PspA/Snf7 superfamily to the last universal common ancestor (LUCA)

We explored the evolution of the PspA-Snf7 superfamily using an MSA of comprehensively selected PspA/Snf7 homologs from distinct clades across the tree of life (49, 50) and different domain architectures (**Fig. 2**). The MSA revealed a unique insertion of heptad repeats in actinobacterial PspA sequences, likely a lineage-specific adaptation. A few cyanobacterial PspA homologs similar to the ancestral plant variant Vipp1 also contained a C-terminal extension (16, 32, 72). A maximum-likelihood phylogenetic tree was generated using this MSA (**Fig. 2**; <u>web app</u>), which showed clear clustering into i) easily distinguishable clades of homologs from Actinobacteria, Firmicutes, and Proteobacteria; ii) the eukaryotic Vipp1 clade grouped with cyanobacterial homologs indicating plastid origins; and iii) Snf7 domain-containing homologs from archaea and eukaryotes forming a well-defined, separate branch. Together, these observations support that the PspA/Snf7 superfamily is ubiquitous across the superkingdoms/domains of life and originated from a PspA-like gene present in the last universal common ancestor (LUCA), as independently corroborated by recent studies (19, 20).

### PspA: Novel architectures and neighborhoods

We used our comprehensive, domain-level search to delineate the domain architectures and genomic contexts of the PspA homologs in organisms from diverse clades.

#### Domain Architectures

Our searches revealed that the underlying domain architectures of most PspA homologs (>98%; ∼2500 homologs) show little variation (<u>web app</u>). In most lineages, PspA homologs contain only the characteristic PspA_IM30 domain (**Fig. 3**; **Table S3;** <u>web app</u>). The remaining <2% of homologs contain PspA_IM30 repeats in tandem or fusions with other domains, including NlpC/P60 and a novel PspA-associated (PspAA) domain (**Fig. 3**; **Text S2.1**). The PspA_IM30–NlpC/P60 hydrolase fusion is predicted to catalyze the modification of phosphatidylcholine, likely altering membrane composition (**Fig. 3**; **Table S3**; <u>AFZ52345.1</u>) (73). Similarly, Snf7 showed minimal variation in domain architecture, with occasional fusions (<5%) in eukaryotes, with some potentially attributable to genome annotation errors. Some Actinobacteria have an Snf7 homolog fused to an RND-family lipid/fatty acid transporter, flanked by two genes encoding a *Mycobacterium*-specific TM protein with a C-terminal cysteine-rich domain (74).

**Figure 3.**
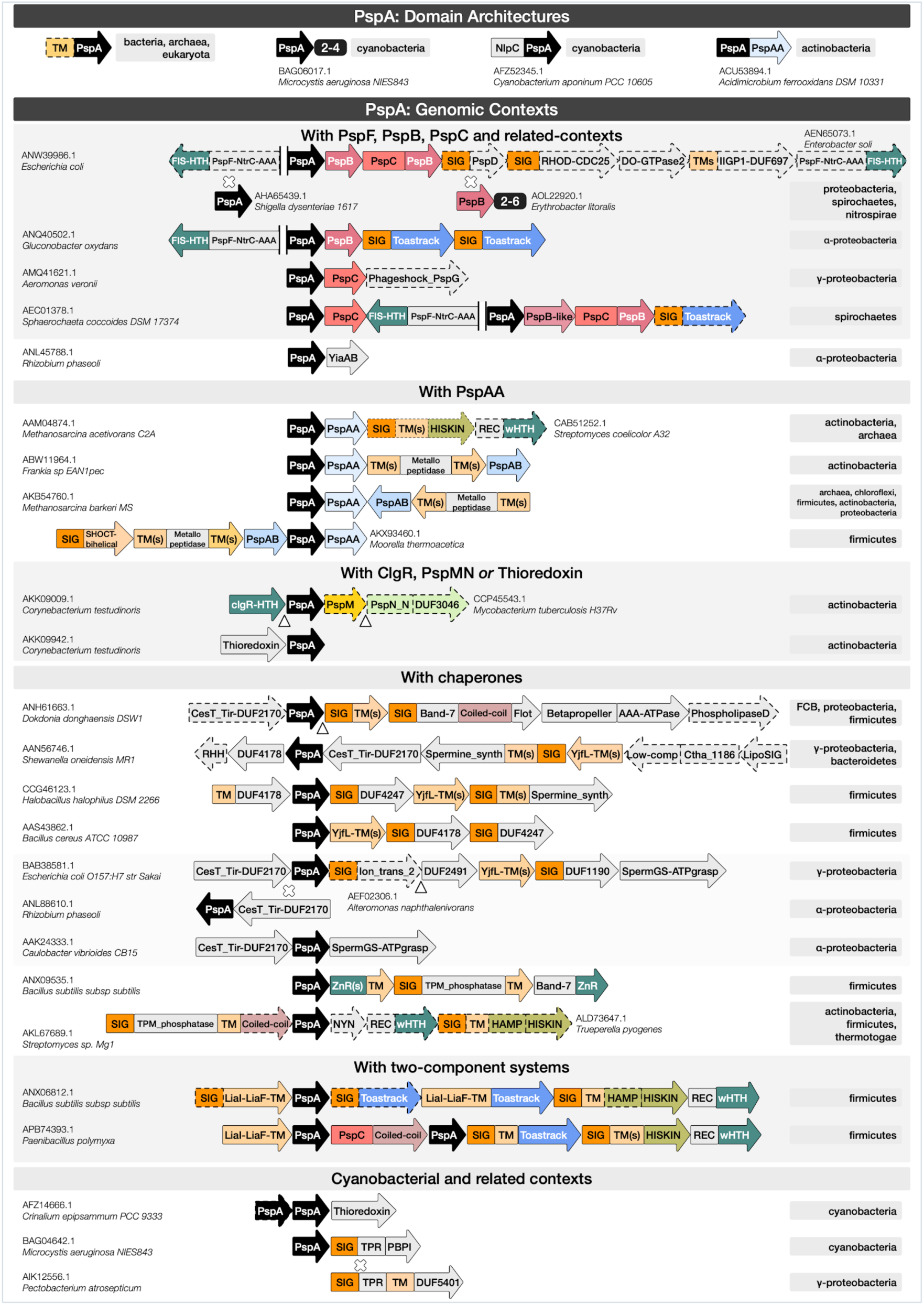
PspA domain architectures and genomic contexts. The first row contains domain architectures of the most prevalent homologs of PspA (as in **1A**; indicated in ***black*** throughout). The remaining rows show the predominant genomic contexts of PspA homologs across multiple bacterial and archaeal lineages (identified by neighborhood searches ±7 genes flanking each homolog; see *Methods*). Representative neighborhoods with archetypal lineages and archetypal example proteins (with accession numbers and species) are shown. The PspA contexts are grouped by neighboring domains such as (a) PspF, PspB/PspC; (b) PspAA, PspAB; (c) ClgR, PspM/PspN, Thioredoxin (**Fig. S1**); (d) chaperones such as Band-7 proteins like flotillin, CesT_Tir, TPM_phosphatase, ZnR, SpermGS-ATPgrasp, and Spermine synthase (**Table S2**); (e) two-component systems such as the Lia system and Toastrack, and other novel genomic contexts; and (f) cyanobacterial variations. <u>Key</u>: *rectangles*, domains; *arrowheads*, the direction of transcription; *domains enclosed by dotted lines*, absent in the genomic contexts in certain species; *white cross*, substitution with protein(s) mentioned just below; *white triangle*, insertion with one or more proteins; *two vertical lines* ‘||’, indicate a change in the direction of transcription; *small black boxes,* domain repeats co-occurring within a protein/context. Archetypal accession numbers and species are provided mostly on the left. Archetypal lineages are indicated in grey on the right for each of the domain architectures and genomic contexts. Different domains are indicated by the same color scheme as in Figure 1. Also, similar domains are given similar hues. For example, membrane proteins are in different shades of orange (SIG, predicted signal peptide, dark orange, PspC, orange, other transmembrane domain, light orange); transcription factors/regulators (including HTH, helix-turn-helix domain) are in teal; DUFs, Domains of Unknown Function, and other domains are in grey. Further details of the domain architectures and gene neighborhoods shown are described in the text and **Table S3**, and the full list of PspA homologs, their domain architectures, genomic contexts, and lineages are shown in the <u>web app</u> (under the ‘Data,’ ‘Domain architectures,’ and ‘Genomic contexts’ tabs).

##### Paralogs

We found that genomes encoding multiple copies of PspA do not maintain domain architecture and genomic context among paralogs (**Fig. 3**; **Table S3**; ‘Phylogeny –> Paralog’ tab, <u>web app</u>). In cyanobacteria, dyads/triads of PspA paralogs occur as adjacent, occasionally fused genes (**Fig. 3**; **Table S3**; *e.g.,* <u>BAG06015.1</u>). Consistent with earlier reports (25, 27), we observed that these neighboring PspA genes are part of two distinct clusters of homologs, one resembling the bacterial PspA and the other the eukaryotic (plastid) Vipp1 (**Fig. 3**). Snf7 gene clusters also appear occasionally in archaeal and eukaryotic species (**Table S3**; *e.g.* <u>CBY21170.1</u>). This suggests tandem gene duplication and may relate to the role of these clusters in membrane stabilization as oligomeric complexes, as in the eukaryotic ESCRT systems (75). Further expansion on PspA/Snf7 paralogs and their likely evolution, including inference of gene duplication *vs.* horizontal gene transfer from domain architectures and genomic contexts, is available in the <u>web app</u>. Occasionally, PspA and Snf7 co-occur in conserved gene neighborhoods (**Fig. 3**; **Table S3**) such as in Deltaproteobacteria (<u>AKJ06548.1</u>); in Bacteroidetes (<u>CBH24266.1</u>), these neighborhoods also contain a third gene encoding a coiled-coil protein with a structure reminiscent of the PspA coiled-coils (**Table S3**). These functional considerations are consistent with recent structural and phylogenetic studies indicating that Vipp1, PspA, and Snf7 proteins have arisen from a common ancestor and play similar roles (19, 20).

#### Novel variations of classical Psp genomic contexts

We next explored the gene-neighborhood themes for each Psp member (**Fig. 3**; **Table S3**). Major themes representing variations of classical Psp operons are summarized below. For the full list of genomic contexts and phyletic spreads, see <u>web app</u>: ‘Data,’ ‘Genomic Contexts’ tabs.

##### Diversity of the pspFABC operon

In addition to Proteobacteria, the classic *pspABC* occurs in nitrospirae and some spirochaetes (**Fig. 3**; **Table S3**). There are also variations to this theme (<u>web app</u>): 1) PspC is fused to a divergent C-terminal PspB in addition to a single PspB in the operon (**Fig. 3**; **Table S3**; *e.g.,* <u>ANW39986.1</u>); 2) PspB duplicates appear in the operon (**Fig. 3**; **Table S3**; *e.g.,* <u>AOL22920.1</u>); 3) PspD occurs along with this operon only in Gammaproteobacteria (**Fig. 3**; **Table S3**; *e.g.,* <u>ANW39986.1</u>). Key variations in the transcription regulator PspF (NtrC-AAA and HTH) include: 1) an operon with additional genes for PspB and PspC fusions and Toastrack-containing proteins in a few spirochaetes; 2) operons in Gammaproteobacteria with genes encoding NtrC-like transcription factors with N-terminal ligand-binding ACT and PAS domains (**Fig. 3**; **Table S3**; *e.g.,* <u>AEN65073.1</u>) (76), and a further protein of the DO-GTPase family, predicted to play a role in membrane-related stresses (77, 78), which occasionally feature an additional PspF; and 3) in some Alphaproteobacteria, the PspC in PspFABC has been replaced by multiple Toastrack-containing proteins (**Fig. 3**; **Table S3**; *e.g.,* <u>ANQ40502.1</u>). Thus, classic *pspABC* presents variation in structure and encoded function.

##### Associations with Vps4 and other classical AAA*+*-ATPases

The core of an ESCRT complex in archaea is defined by the co-occurrence of one or more copies of Snf7 (*e.g.,* <u>OLS27540.1</u>; [**Table S3**]) and a gene encoding a **VPS4-like AAA+-ATPase** (with an N-terminal <u>MIT</u> and C-terminal oligomerization domain [**Table S2**]) (79). In our analysis, we observed diversity in *vps4* gene neighborhoods (**Text S2.2**). ESCRT complexes are well-studied in eukaryotes, but less so in archaea, where they were originally associated with cell division and named for this role (Cdv system) (80). Crenarchaeota and Thaumarchaeota contain *cdvABC*. *cdvA* has no eukaryotic homolog, while *cdvB* and *cdvC* encode homologs of ESCRT-III proteins and Vps4, respectively. Recent studies have shown that eukaryotes originate within the archaeal clade of Asgardarchaeota (80). Consistent with this, Asgardarchaeota show a conserved gene neighborhood coupling Snf7, a VPS4-like AAA+ ATPase gene, and a gene coding for ESCRT-II wHTH-domain (81) corresponding to the cognate eukaryotic ESCRT complex. These operons may be further extended with additional copies of Snf7 genes and other genes coding for a TM protein and an ABC ATPase. Additionally, examples of transfers of VPS4-like AAA+-ATPase from archaea to several bacterial lineages were observed (**Table S3**; *e.g.,* <u>ACB74714.1</u>). In these cases, *snf7* is displaced by an unrelated gene containing TPR repeat and 6TM, suggesting membrane proximity. The bacterial PspA (**Table S3**; *e.g.* <u>AEY64321.1</u>) may occur with a distinct protein containing two AAA+-ATPase domains – with the N-terminal copy inactive – in some bacterial clades (**Text S2.2**). Both PspA and the membrane-associated Snf7, along with the AAA+-ATPase, may occur in longer operons with other genes encoding an ABC-ATPase, permease, and a solute- and peptidoglycan-binding protein with PBPB-OmpA domains (**Table S3**; *e.g.,* <u>OGG56892.1)</u>. These associations between particular bacterial PspA and Snf7 clade members with AAA+-ATPases suggests a role in ATP-dependent membrane remodeling (**Fig. 3**; **Table S3**), comparable to Snf7 and VPS4 in the ESCRT-III-mediated membrane remodeling in the archaeo-eukaryotic lineage (19, 20, 61, 68, 69, 71, 75). The additional transporter components seen in some of these systems suggest a link between this membrane remodeling and solute transport. Here the OmpA domain could interact with peptidoglycan, while the PBPB domain binds the solute trafficked by the linked ABC transporter.

##### Operons with CesT/Tir-like chaperones and Band-7 domain proteins

We discovered two novel overlapping genomic associations across various bacteria linking PspA with the Band-7 domain and CesT/Tir-like chaperone domain proteins (**Fig. 3**; **Tables S2** and **S3**). Band-7 has been implicated in macromolecular complex assembly, playing a chaperone-like role in binding peptides (82, 83), and in both membrane-localization and membrane dynamics (84). CesT/Tir chaperone domains have been shown to mediate protein-protein interactions in the assembly and dynamics of the Type-III secretion systems of proteobacteria (85). Using profile-profile searches, we identified a previously uncharacterized protein encoded by genes linked to the yjfJ family of *pspA* genes (*e.g.,* <u>ANH61663.1</u>) as a novel member of the CesT/Tir superfamily (**Fig. 3**; **Tables S1** and **S2**; *e.g.,* <u>ANH61662.1</u>). We also observed that the proteobacterial proteins in the neighborhood of *pspA* (**Fig. 3**; *e.g.,* <u>BAB38581.1</u>), typified by YjfI from *E. coli* (<u>DUF2170</u>; **Table S2**; *e.g.,* BAB38580.1), contained a CesT/Tir superfamily domain (86).

The dyad encoding CesT/Tir–PspA forms the core of one class of conserved gene neighborhood (**Table S2**). We observed it as either a standalone unit or within larger operons, such as one group combining CesT/Tir–PspA (**Fig. 3**; **Table S3**; <u>ANH61663.1</u>) with i) a gene encoding a membrane-associated protein with the domain architecture TM+Band-7+coiled-coil+flotillin, ii) a novel AAA+-ATPase fused to N-terminal coiled-coil and β-propeller repeats, and iii) a 3TM domain protein prototyped by *E. coli* YqiJ and *B. subtilis* YuaF (previously implicated in resistance to cell wall-targeting antibiotics) (87, 88). In some Proteobacteria, this operon also contains a phospholipase D (HKD) superfamily hydrolase (**Fig. 3**). Related abbreviated operons coding only for the Band-7 and flotillin-domain-containing protein, the YqiJ/YuaF TM protein, and in some cases, the aforementioned AAA+-ATPase protein is more widely distributed across bacteria and archaea. These might function with PspA homologs encoded elsewhere in the genome.

The second major group incorporates the dyad into a 6–7 gene operon (**Fig. 3**; **Table S3**; *e.g.,* <u>AAN56746.1</u>) featuring a gene encoding a spermine/spermidine-like synthase domain (89) fused to an N-terminal 7TM transporter-like domain (**Fig. 3**; **Tables S2** and **S3**; *e.g.,* <u>AAN56744.1</u>). This operon encodes three additional uncharacterized proteins, including the YjfL family 8TM protein (<u>DUF350</u>), a novel lipoprotein (Ctha_1186 domain), and a β-strand rich protein that we predict to be intracellularly localized (**Fig. 3**; **Tables S3** and **S4;** <u>DUF4178</u>; *e.g.,* <u>AAN56747.1</u>). In some cases, the last gene in this operon codes for a ribbon-helix-helix (RHH) domain transcription factor (**Fig. 3**; **Tables S2** and **S3**).

The third group combines the CesT/Tir–PspA dyad (<u>AMJ95269.1</u>) with polyamine metabolism genes encoding an ATP-grasp peptide ligase (<u>AMJ95273.1</u>) related to the glutathionyl spermidine synthetase (**Fig. 3**; **Table S3**). This association was also recently noticed in a study of AdoMet decarboxylase gene linkages (90). The operon also encodes a potassium channel with intracellular-ligand-sensing, a YjfL family 4TM protein (**Tables S2** and **S3**; <u>DUF350</u>), a metal-chelating lipoprotein (<u>DUF1190</u>), and another uncharacterized protein with predicted enzymatic activity (<u>DUF2491</u>).

Alternately, two variants of these operons lack the CesT/Tir–PspA dyad: one where its replacement is a gene encoding a distinct protein occurring as either a secreted or lipoprotein version (<u>DUF4247</u>, *e.g.,* <u>CCP45395.1</u>) and a predicted beta-sandwich enzymatic domain with two highly conserved histidines, as well as a conserved aspartic acid and glutamic acid (<u>DUF2617</u>, *e.g.,* <u>CCP45394.1</u>). In the second variant, the dyad is replaced by a protein containing a Band-7 domain fused to C-terminal 2TM and SHOCT domains (see below). These operons encode two further polyamine metabolism genes, an AdoMet decarboxylase, and a polyamine oxidase.

Lastly, the *Bacillus subtilis ydjFGHL* operon typifies a coupling in Firmicutes of *pspA* (**Table S1**; <u>ANX09535.1</u>) with Band-7 and C-terminal Zinc Ribbon (ZnR) domains in YdjL, and two ZnRs followed by an uncharacterized domain related to YpeB featuring an NTF2-like fold and a TM segment in protein YdjG. The conserved partner of PspA in these systems is the membrane-associated YdjH, which is characterized by a so-called “**TPM phosphatase domain**” followed by a low-complexity segment or a long coiled-coil (**Fig. 3**; **Table S3**) (91). Competing hypotheses suggest that this domain either encodes a generic phosphatase based on its plant homolog (91) or is involved in membrane-associated photosystem II repair and respiratory complex III assembly (92, 93). Thus, TPM association with PspA parallels the associations with the chaperone CesT/Tir and Band-7 domains involved in macromolecular assembly. Our result supports the hypothesis that the CesT/Tir, Band-7, and TPM domains associate with PspA to serve a chaperone-like role in the assembly of membrane-linked protein complexes.

##### Membrane dynamics with Chaperone-like domains

The contextual association of PspA with AAA+-ATPase, two distinct families of CesT/Tir type chaperones, Band-7 and TPM domains further supports functional roles of PspA homologs in recruiting macromolecular assembly machinery to the inner leaf of the cell membrane. Polyamines like spermine/spermidine have been implicated in membrane stability in a concentration-dependent manner (94). The repeated coupling of polyamine metabolism genes we identified in PspA-CesT/Tir operons implies that a PspA-based system of membrane dynamics monitors polyamine concentration or aminopropylation of other substrates (90) to alter membrane structure and membrane-associated protein complexes. We propose that these systems link extracellular stress sensing with intracellular membrane dynamics based on the frequent coupling with cell-surface lipoproteins. In particular, flotillin domain-containing proteins are likely to be recruited to lipid subdomains with special components (*e.g.,* cardiolipin-rich inner membrane) to interface with PspA (88).

##### The PspAA domain

Our *pspA* gene neighborhood analysis identified a unique partner in archaeal and bacterial phyla: a protein containing a novel domain with 3 helices and a 4-stranded β-sheet occurring in a two-gene cluster with *pspA* (**Fig. 3**; **Table S3**; *e.g.,* <u>AAM04874.1</u>). This domain often occurs independently but is occasionally fused to an N-terminal PspA in Actinobacteria and Chloroflexi (**Fig. 3**; **Table S3**; *e.g.,* <u>ACU53894.1</u>). This **PspA-associated domain A (PspAA)** (**Fig. 3**; **Table S2**; **Text S1.3**) occurs in a PspA–PspAA dyad in a two-component system (*e.g.*, <u>CAB51252.1</u>), and in another system consisting of a membrane-associated metallopeptidase, another novel domain **PspA-associated domain B (PspAB)** (*e.g.*, <u>AAZ55047.1</u>), and an occasional third gene encoding a SHOCT-like domain (**Fig. 3**; **Tables S2** and **S3**; <u>web app</u>).

##### PspA with PspM (ClgRPspAMN) or Thioredoxin

Certain *pspA* homologs may occur in Cyanobacteria and Actinobacteria as an operonic dyad with a gene encoding an active thioredoxin domain-containing protein (15). PspA in this dyad is typified by corynebacterial RsmP (*e.g.,* <u>AKK09942.1</u>; **Fig. 3**; **Tables S2** and **S3**; **Text S2.3**), which is predominantly found in rod-shaped Actinobacteria and regulates cell shape via phosphorylation (95). These PspA homologs either occur with thioredoxin (*e.g.,* <u>AKK09942.1</u>) or with a ClgR-like cHTH and PspM genes (*e.g.,* <u>CCP45543.1</u>; **Fig. 3, Fig. S1, Table S3**). Corynebacterium spp. Contain two distinct PspA paralogs each displaying one of the above contextual associations (‘Paralog’ tab, <u>web app</u>). The same thioredoxin family occurs with a different PspA clade (typically two copies) in Cyanobacteria (*e.g.,* <u>AFZ14666.1</u>; **Fig. 3**; **Table S3**). The association of the ClgR-PspM and thioredoxin and their mutual exclusion in the operon suggest that they act as alternative regulators of the PspA homologs — repressors in the case of the ClgR, and a potential redox regulator in the case of thioredoxin (**Fig. S1**; **Text S2.3**).

#### Operons with two-component systems

Multiple two-component systems associate with the PSP system. For instance, the *liaIHGFSR* operon from *B. subtilis* has been studied in the context of lantibiotic resistance (*e.g.*, <u>ANX06812.1</u>; **Fig. 3**; **Table S3**) (11). In a few paenibacilli, genes encoding an additional PspA and PspC with a C-terminal coiled-coil have been inserted into a *liaIHGFSR*-like operon (*e.g.,* <u>APB74393.1</u>; **Fig. 3**; **Table S3**). In Actinobacteria (*e.g.*, <u>CAB51252.1</u>; **Fig. 3**; **Table S3**), the PspA–PspAA dyad occurs with a firmicute-like two-component system.

##### Classic two-component transcriptional signaling system

We identified two-component signaling system operons linked to either *pspA* or *PspC* in Firmicutes, Actinobacteria, and other clades (**Fig. 3**; <u>web app</u>). In these operons, a membrane-bound histidine kinase (HISKIN) communicates an external stress signal to the response regulator (a receiver domain fused to a HTH domain) protein, triggering a transcriptional response. It is very likely that PspA, LiaI-LiaF-TM, and Toastrack tie into a two-component system regulating membrane integrity and permeability in response to the stress signal. In Actinobacteria, where the HISKIN is fused to PspC, the signal is presumably sensed by PspC. Moreover, when *pspC/Toastrack* operons with two-component systems lack *pspA* in their immediate operonic neighborhood, they might recruit PspA proteins from elsewhere in the genome to express stress response functions.

### PspA-free variations of domain architectures and gene-neighborhood

We next investigated domain architectures and genomic contexts involving PSP components that lack PspA, including those with the PspC and Toastrack domains.

#### PspC domain architectures and gene-neighborhood

**PspC** is present in most bacteria, some archaeal clades of Euryarchaeota, and a few Asgardarchaeota (<u>web app)</u>. Some orthologs of PspC are fused to an N-terminal ZnR domain (*e.g.,* <u>ABC83427.1</u>, **Table S4**), while most other homologs occur either in predicted operons or fusions with multi-TM domains, such as the LiaI-LiaF-TM and PspB (**Fig. 4**; **Table S4**). PspC also fuses to diverse signaling domains such as i) HISKIN from two-component systems (see above); ii) a novel signaling domain we term the <u>H</u>TH-<u>a</u>ssociated <u>α</u>-helical <u>s</u>ignaling domain (**HAAS)**, which overlaps partly with the Pfam model for <u>DUF1700</u> (**Table S2**), and iii) the Toastrack domain (**Table S1**).

**Figure 4.**
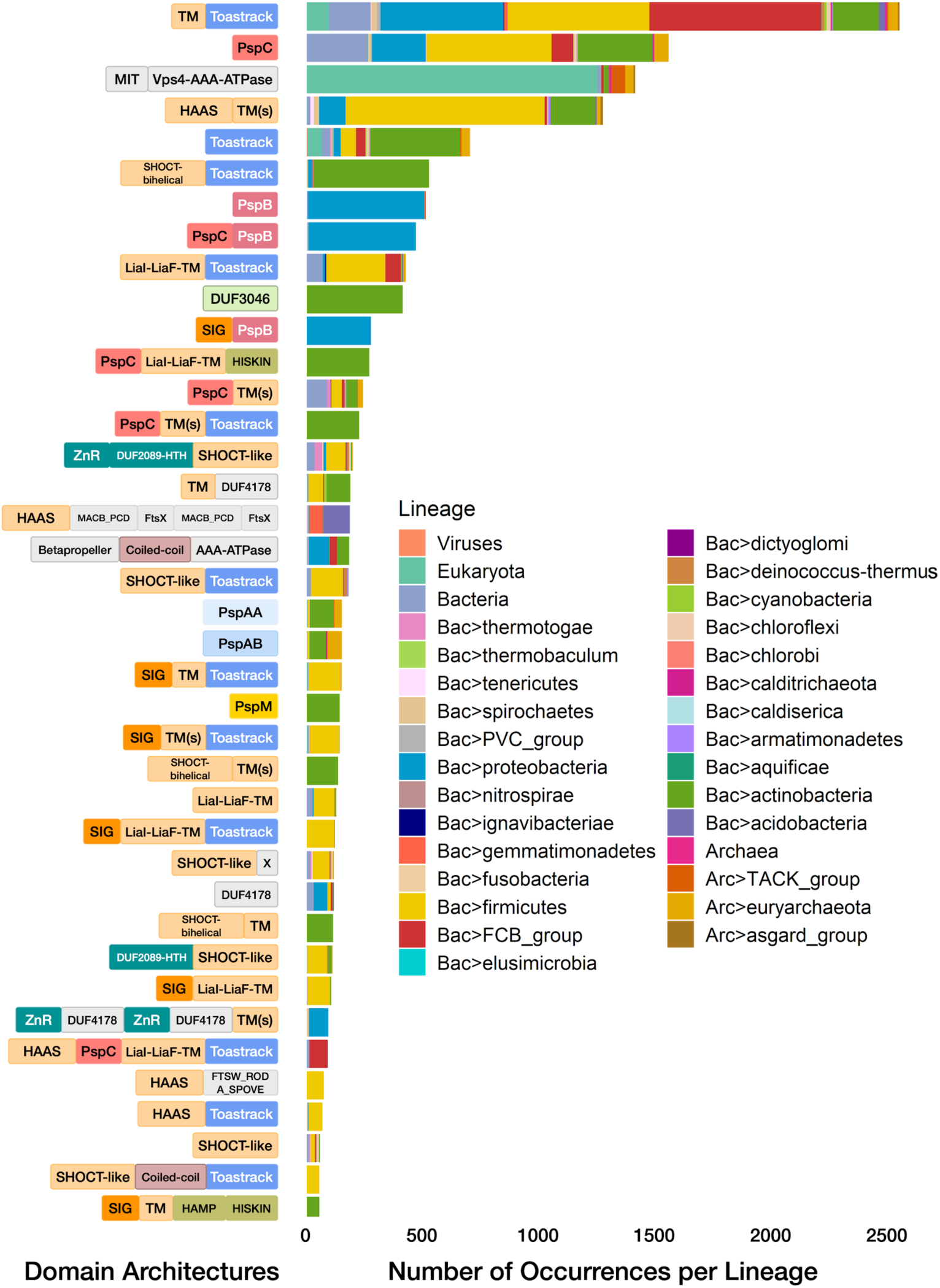
Lineage spread of PspA-free domain architectures. The domain architectures of the most prevalent homologs of PspA partner domains (frequency of occurrence >50 across lineages), including classical (Toastrack, LiaI-LiaF-TM, PspBC, PspMN, DUF3046) and other novel neighbors (PspAA, PspAB, HAAS, SHOCT-bihelical, SHOCT-like, AAA+-ATPase domains) are illustrated on the left. Broad distributions of domains across the tree of life are indicated in Fig. 1C. The phyletic spread of the underlying domain architectures is shown here along with their relative frequencies as a stacked barplot for all superkingdoms/domains of life (sub-lineages or phyla are sorted; *e.g.,* all bacterial lineages appear together). Further details of the domain architectures of all PspA partner domain homologs and their phyletic spreads are shown in the <u>web app</u>, with representatives shown in **Table S4**.

Moreover, Actinobacteria contain a Lia-like system without PspA. The HISKIN domain in these systems is fused to N-terminal PspC and LiaF-LiaI-TM domains (*e.g.,* <u>AIJ13865.1</u>) and accompanied by a REC–HTH-containing protein (**Fig. 5**; **Table S4**). This core shares a promoter region with a second gene on the opposite strand containing a PspC domain that is fused to additional TMs and, in some cases, a Toastrack domain (*e.g.,* <u>AIJ13866.1</u>; **Fig. 5**; **Table S4**). These associations strongly imply that PspC acts as a sensor domain to detect and communicate a membrane-proximal signal.

**Figure 5.**
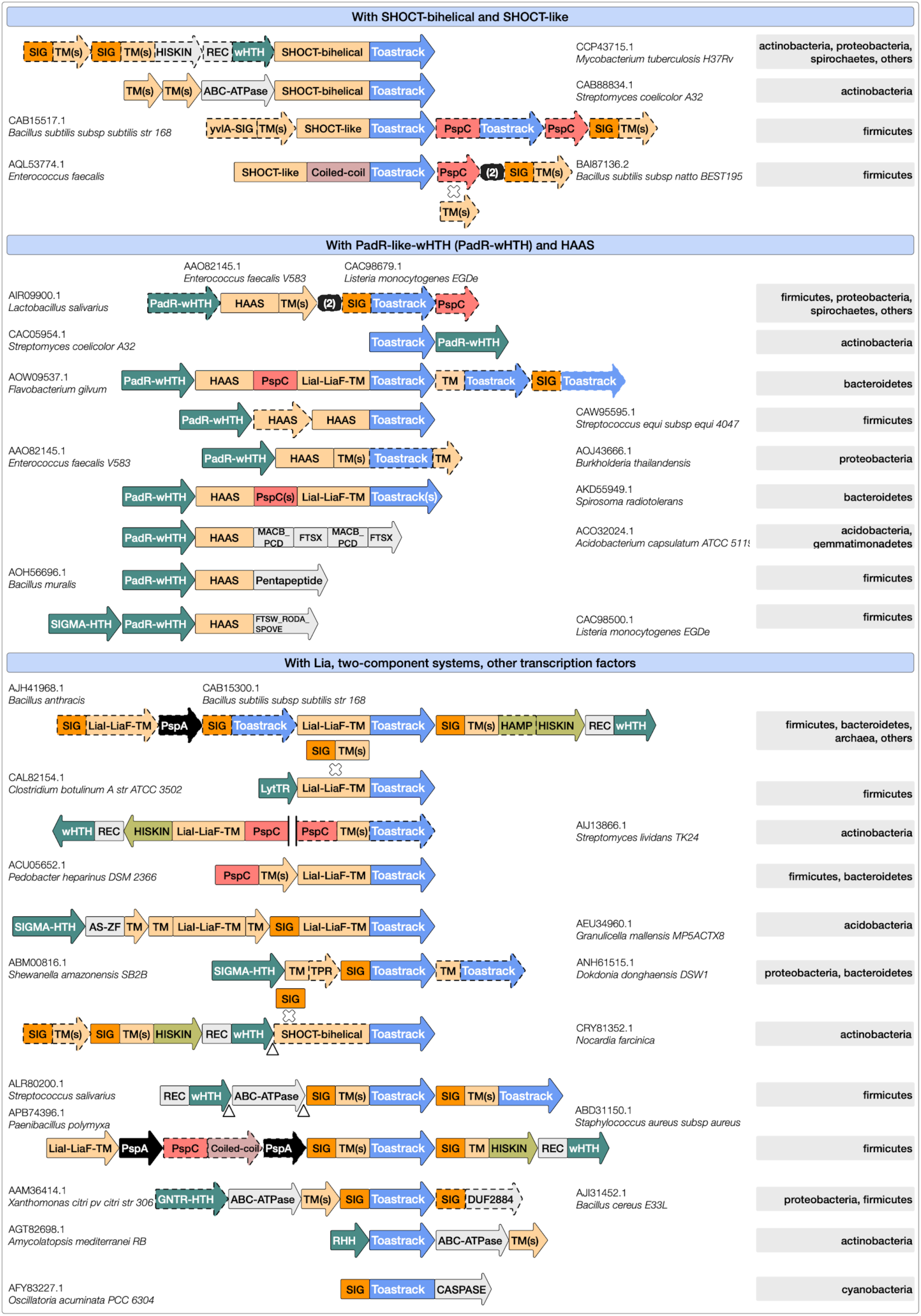
The genomic contexts housing all PSP cognate partner domain homologs. The genomic contexts and key lineage memberships are presented using the same schematic as in Figure 3. The focus is on PSP partner domains such as Toastrack (blue), PspC, LiaI-LiaF-TM, HAAS, SHOCT-bi-helical (in shades of orange since they are transmembrane domains), and the various genomic neighborhoods with SHOCT-like proteins, transcription regulators (*e.g.,* PadR-wHTH, SIGMA-HTH, GNTR-HTH), and two-component systems (**Table S2**). Further details of the genomic contexts of all PspA-free partner domain homologs and their phyletic spreads are in the <u>web app</u>, and representatives indicated in the figure are shown in **Table S4**.

#### Contextual associations of the Toastrack Domain

The **Toastrack** domain repeatedly emerges as a partner to the PSP system. As noted, genes encoding Toastrack-containing protein occur concurrently with *pspA* in several conserved neighborhoods (**Fig. 4**; **Table S4**) and are fused to *pspC* in *pspA*-free settings. We found that the Toastrack domains tend to be C-terminal to various TM domains such as PspC, LiaI-LiaF-TM, and other multi-TM/single-TM domains unique to these systems (**Fig 4**; **Text S2.4**). These fusions, along with the lack of TM or extracytoplasmic regions within Toastrack (**Table S1**; **Figs. 1** and **4**), strongly suggest that Toastrack is an intracellular domain. Further, the N-terminal regions of several Toastrack architectures contain at least two divergent homologs of the bi-helical SHOCT domain (*e.g.,* <u>CAB15517.1</u>; **Figs. 4** and **5**; **Tables S1** and **S4**), which we call **SHOCT-like** domains, which are partly detected by the Pfam **<u>DUF1707</u>** model (**Text S2.4**) (96). We observed these SHOCT-like domains at both N- and C-termini of various globular domains, including Toastrack. The globular domains fused to the SHOCT and related domains identified in this work may occur independently of them in other contexts. In most cases, the domains are neither fused to signal peptides nor to TM segments to indicate a potential extra-cellular location. Moreover, their known/predicted activities are meaningful only in intracellular contexts. Therefore, we predict that this SHOCT-like domain may play a role in anchoring disparate domains to the inner leaf of the membrane.

The DNA-binding domain **LytTR** may also be encoded by operons carrying Toastrack fused to LiaI-LiaF-TM in Firmicutes (*e.g.,* <u>CAL82154.1</u>; **Fig. 5**; **Table S4**). The Toastrack-containing protein (*e.g.,* <u>CAB15517.1</u>) occurs in an operon with a *pspC* homolog (<u>CAB15516.1</u>) and has been implicated in the activation of the membrane-associated *liaFSR* operon (**Table S1**) and protection against membrane permeabilization (97–99). We also identified *pspA*-free variants of the classic *lia* operon with Toastrack and a two-component system, typified by *vraT* of *Staphylococcus aureus* (<u>ABD31150.1</u>; **Fig. 5**; **Table S4**). These systems carry LiaI-LiaF-TM and Toastrack with a two-component system (*vraSR*) and are involved in methicillin and cell wall-targeting antibiotic resistance (3, 100–102). A similar organization of LiaI-LiaF-TM and Toastrack containing a protein with a two-component system is found in a few bacterial species (*e.g.,* among Proteobacteria, <u>ABD83157.1</u>, Ignavibacteriae, <u>AFH48155.1</u>; **Table S4**).

We discovered several other conserved gene neighborhoods centered on Toastrack-containing genes likely to define novel functional analogs of the PSP system with potential roles in membrane-linked stress response. One of these, a four-gene context found across diverse bacterial lineages, includes a previously uncharacterized protein similar to the Pfam model <u>DUF2089</u>. The Pfam model **DUF2089** can be divided into an N-terminal ZnR, central HTH, and C-terminal SHOCT-like domains (<u>ADE70705.1</u>), variants of which appear with LiaI-LiaF-TM, B-box, or PspC domains (**Figs. 4** and **5**; **Tables S2** and **S4**; **Text S2.4**). Because of these shared operonic associations, these Toastrack contexts may function similarly to the classical *lia* operon in transducing membrane-associated signals to a transcriptional output affecting a wide range of genes, including through an operonically linked sigma factor.

Similarly, we observed operons coupling a membrane-anchored Toastrack-containing protein with an ABC-ATPase, a permease subunit, a **GNTR-HTH** transcription factor with distinct C-terminal α-helical domain (**Fig. 5**; **Tables S2** and **S4**; **Text S2.4**), and a <u>DUF2884</u> membrane-associated lipoprotein (*e.g.,* <u>AJI31452.1</u>). The latter might function as an extracellular solute-binding partner for the ABC-ATPase and permease components. We found a similar operon in Actinobacteria with the ribbon-helix-helix (**RHH**) domain transcription factor replacing **GNTR-HTH** (**Table S2**; **Text S2.4**). In some Actinobacteria, the Toastrack domain is fused to a SHOCT-like domain or occurs in a transport operon, likely coupling Toastrack to transcriptional regulation, membrane-proximal signal sensing, and transport (**Fig. 5**; **Table S4**).

Based on these observations, we propose that the two extended β-sheets of the Toastrack domain (formed by the β-helical repeats) facilitate the assembly of sub-membrane signaling complexes by providing two expansive surfaces amenable to protein-protein interactions near the membrane. Protein docking to the Toastrack domain may also activate transcription via fused or associated HTH or LytTR domains, associated two-component systems, or the HAAS-HTH system.

##### The HAAS-PadR-like wHTH systems

We discovered a novel signaling system that frequently co-occurs with PspC and Toastrack domains in gene neighborhoods that do not contain *pspA* (**Figs. 4** and **5**). This system consists of two components, the HAAS domain and the PadR-like wHTH (**Table S2**). The HAAS domain is partly similar to the Yip1 protein family (Pfam models <u>DUF1700</u> and <u>DUF1129</u>; Pfam clan: <u>Yip1</u>). An HHpred search further unifies this model with the Pfam profile <u>DUF1048</u> (PDB: <u>2O3L</u>). We therefore refer to them collectively as the **HAAS** superfamily. The core HAAS fold has three consecutive α-helices, with the third helix displaying an unusual kink corresponding to a conserved GxP motif, predicted to form part of a conserved groove that mediates protein-protein interactions. The HAAS domain always occurs in gene neighborhoods coupled with a protein containing a standalone PadR-like wHTH DNA-binding domain (**Fig. 5**). This co-migration suggests that HAAS transduces a signal from domains fused to this wHTH transcription factor.

When coupled with PspC, the HAAS domain shows three broad themes: 1) it is found as part of a multidomain TM protein with additional PspC, LiaI-LiaF-TM, and Toastrack domains; 2) it is fused directly to a Toastrack domain in a gene neighborhood that also codes for a PspC domain protein, occasionally fused to other TM domains; 3) it constitutes a part of a TM protein fused to conserved multi-TM domains other than the LiaI-LiaF-TM or PspC (*e.g.,* VanZ) (**Figs. 4** and **5**; **Table S4**). These gene neighborhoods code for standalone *pspC* genes. Additionally, the HAAS domains occur in PspC-independent contexts, but are typically coupled to the N-terminus of other multi-TM or intracellular sensory domains. We found conserved fusions to at least 5 distinct multi-TM domains (**Figs. 4** and **5**; **Table S4**), such as 1) the FtsX-like TM domains with extracellular solute-binding MacB/PCD domains (*e.g.,* <u>ACO32024.1</u>); 2) FtsW/RodA/SpoVE family of TM domains (*e.g.,* <u>CAC98500.1</u>) (103); 3) various uncharacterized 6TM, 4TM, and 2TM domains. In addition to Toastrack, the HAAS domain may combine with other intracellular domains such as the pentapeptide repeat domains mostly in Firmicutes (*e.g.,* <u>AOH56696.1</u>; **Fig. 5**; **Table S4**). The organisms in which these operons occur have PSP components elsewhere in the genome. However, these proteins might be recruited to the stress response system, as demonstrated by the effects of PadR-like wHTH deletion studies in the *Listeria* Lia systems (103).

In bacteria lacking the typical *lia* operon (carrying the LiaRS-like two-component system), we noted that the HAAS–PadR-like-wHTH dyad replaces the histidine kinase-receiver domain transcription factor pair. Therefore, we propose that the HAAS-PadR dyad constitutes an alternative to the classical two-component systems. Here, the HAAS domain is proposed to undergo a conformational change in response to membrane-linked or intracellular signals detected by the sensor domains (*e.g.,* LiaI-LiaF-TM or PspC) that are fused to or co-occur with the dyad. This conformational change is likely transmitted to the associated PadR-like wHTH via the groove of the conserved GxP motif; this change might lead to transcription factor release and a downstream transcriptional response.

### A unified view of PSP partners and evolution

We reconcile our findings on PSP-related genomic contexts and partner domains within the framework of the tree of life to assemble the PSP puzzle. The above results include ∼20,000 homologs of key PSP proteins, ∼200 novel domain architectures (minimum 2 species), and ∼500 genomic contexts (minimum 5 species) (<u>web app</u>). To visualize the broader context of PSP systems across the superkingdoms, we first built a network of domain architectures for the extended PSP system (**Fig. 6A**), with domains as nodes and co-occurrence within a protein as edges. This network consolidates the PSP system with known and novel connections, including those between PspA and PSP partner domains. Next, we summarized the phyletic spread and prevalence of all PSP members and partners in a single heatmap (**Fig. 6B**). This view indicates that, of the PSP components, (i) only PspA/Snf7 is present in all eukaryotes, whereas Toastrack has been transferred from bacteria to certain eukaryotes (*e.g.,* human FAM185A with orthologs in animals and other eukaryotes) ; (ii) PspC and PspA/Snf7 are present in most archaeal lineages, (iii) occasional transfers of Toastrack, LiaI-LiaF-TM, and SHOCT domains are observed from bacteria to Euryarchaeota; (iii) in bacteria, domains such as Toastrack, PspC, PspA, and LiaI-LiaF-TM tend to co-migrate. Finally, using an ‘upset plot,’ we unified: i) the relative occurrence of domain architectures from PSP members and their most common neighbors, ii) combinations of these domain architectures, and iii) their frequencies (**Fig. 6C**). These consolidated visualizations can be explored in our <u>web app</u>. For readers who wish to explore any of these proteins in more depth, the accession numbers from this dataset can also be used for input in our MolEvolvR protein analysis web app (https://jravilab.org/molevolvr) (104).

**Figure 6.**
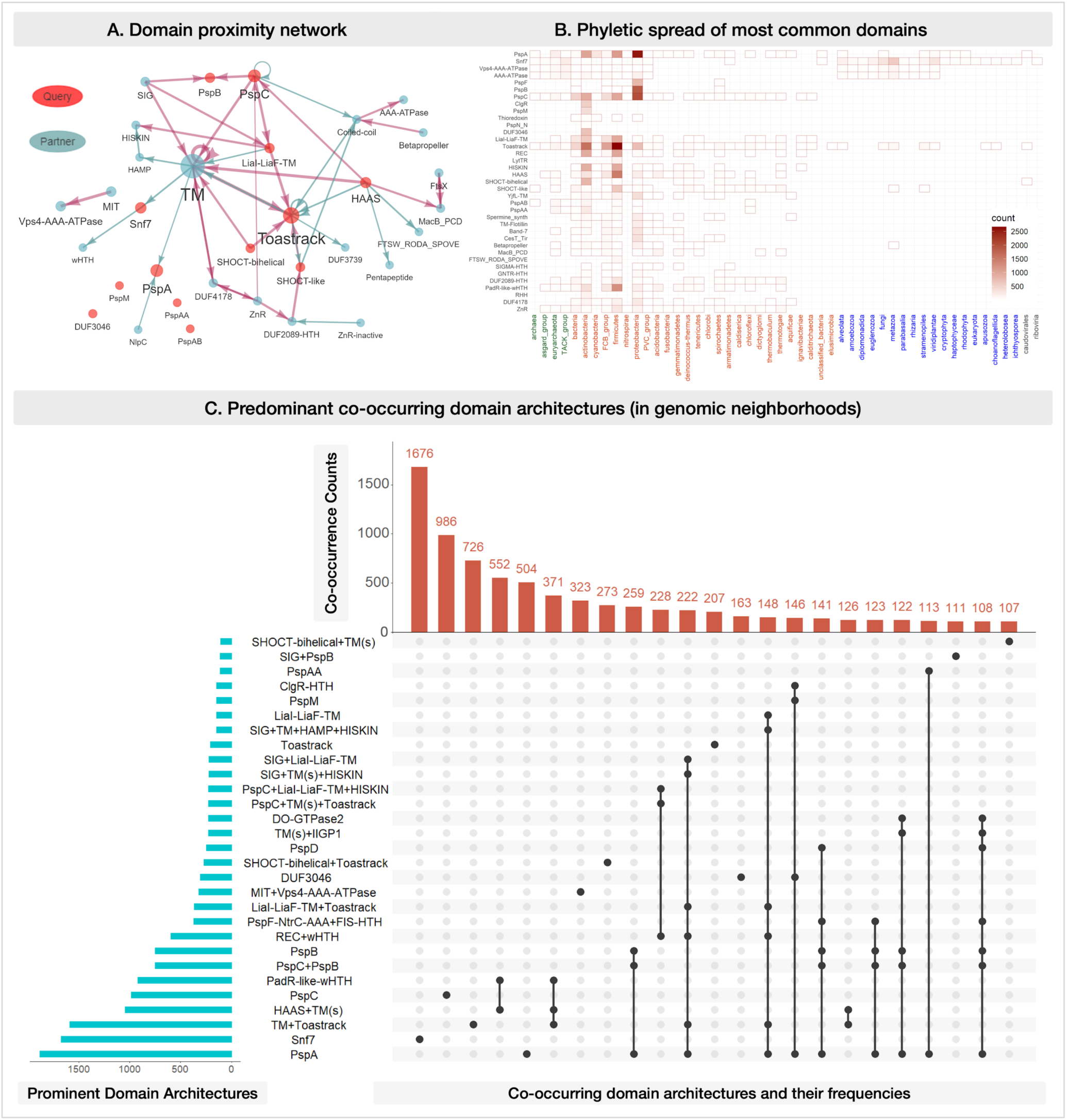
PSP consolidated. **A. Domain proximity network.** The network captures co-occurring domains within the top 97% of the homologs of all the ‘query’ Psp members and their key partner domains (after sorting by decreasing frequency of occurrence). The size of the nodes (domains) and width of edges (co-occurrence of domains within a protein) are proportional to the frequency of their occurrence across homologs. The query domains (original proteins/domains of interest; **Table S1**) and other commonly co-occurring domains (**Table S2**) are ***red*** or ***grey***. *Note*: A few connections may be absent in the displayed network due to low occurrence (fewer than the threshold), *e.g.,* PspA and PspAA, Betapropeller, and AAA-ATPase. The full network, and the domain-centric ones, can be accessed on the <u>web app</u>. **B. Phyletic spread of the most common domains.** The heatmap shows the presence/absence of homologs of PspA and partner domains across lineages. The color gradient represents the number of homologs identified within each lineage (*e.g.,* the darkest shade indicates the highest number of homologs in a particular lineage, white indicates the lowest count, and the absence of a shaded box represents a lack of available data). *Rows*: PSP members and their most frequent partners are queried against all sequenced and completed genomes across Bacteria, Eukaryota, and Archaea. *Columns*: The major archaeal (***green***), bacterial (***orange***), eukaryotic (***blue***), and viral (***grey***) lineages with representative sequenced genomes are considered. Details of all homologs across the tree of life, their domain architectures, genomic contexts, and their lineage distributions are shown in the <u>web app</u> (representatives in **Table S3, S4**). **C. Predominant co-occurring domain architectures in genomic neighborhoods.** UpSet plot of the most common neighboring proteins (genomic contexts >100 occurrences are shown) among all Psp homologs. *Blue histogram*: Distribution of the predominant domain architectures. *Dots and connections*: Combinations in which these domain architectures come together in the genomic neighborhoods. *Red histogram*: Frequency of occurrences of genomic neighborhoods comprising specific combinations of the predominant domain architectures. Phyletic spreads and UpSet plots of the domain architectures and genomic contexts for the homologs of all PSP member proteins are available in the <u>web app</u>.

## Conclusion

Recent work with the bacterial PSP stress response system suggests that while function is maintained across phyla, the proteins, regulatory mechanisms, and membrane stress response mechanisms vary widely among lineages (12, 15, 26, 36). These variations reflect the utilization of a common peripheral inner membrane protein in lineage-specific envelope dynamics and stress responses. In this study, we systematically analyzed PSP systems, their phyletic spread, and evolution (**Figs. 1** and **2**), discovering several novel sequence and structural features (*e.g.,* domain architectures) and conserved gene-neighborhoods of PSP components. We identified both classical and newly-predicted partners (**Figs. 2** and **5**) and established their phyletic distributions across the tree of life (**Figs. 2** and **6**). We also established that PspA/Snf7 traces to the LUCA (**Fig. 2**). Indeed, it has been independently shown that the PspA/Vipp1 ESCRT system dates back to the LUCA (19, 20), further strengthening the evolutionary significance of our finding. This confirms that the LUCA possessed a membrane whose curvature and dynamics were mediated by an ancestral PspA/Snf7-like coiled-coil protein assembled into polymeric structures adjacent to the inner leaf of the membrane. This hypothesis is in line with earlier work showing that LUCA contained some signal-recognition particle GTPases (105). These proteins are unequivocal indicators of the presence of a secretory apparatus translocating proteins across lipid bilayers. Hence, as a corollary they are also indicators of the presence of a membrane. Similarly, the extant components of membrane-linked bioenergetics (F1/F0 ATPases or V-Type ATPases) are also indicative of a membrane in the LUCA (106, 107). Thus, we infer that this ancient machinery was robust to the subsequent lineage-specific changes in membrane chemistry, producing different membrane types (*e.g.,* ether-linked vs. ester-linked) in the archaeal, bacterial, and eukaryotic lineages.

Our novel PSP findings make several new functional predictions: 1) Based on its occurrence with membrane partners (*e.g.,* PspA, LiaI-LiaF-TM, PspC, HAAS, SHOCT-like domains) and transcription regulators or two-component systems (*e.g.,* LiaRS, PadR-like-wHTH), we propose that Toastrack is a membrane-proximal intracellular domain serving as a sub-membrane signaling complex assembly platform; 2) The association of PspA homologs with distinct AAA+-chaperones and a non-ATPase chaperone (*e.g.,* CesT/Tir) or predicted chaperone-like proteins (*e.g.,* Band-7, TPM) suggest that these complexes are involved in the assembly of sub- and trans-membrane complexes in response to specific envelope stresses. These observations imply a general role for analogs of ESCRT-like complexes in these bacterial systems; 3) The association of PspA with various transporters and polyamine metabolism systems indicates that the regulation of membrane structure by PspA is associated with changes in solute concentrations and covalent modifications that may affect membrane stability; 4) The discovery of several new versions of the SHOCT-like domain suggests that these domains mediate membrane localization of disparate catalytic and non-catalytic activities, which may subsequently interface with the PSP system; 5) A diverse array of sub-membrane (*e.g.,* Toastrack), TM, and cell-surface domains (*e.g.,* PspC, LiaI-LiaF-TM) co-occur with two-component or HAAS-PadR-like-wHTH systems, hinting at alternatives to conventional two-component signaling that interface with the PSP system in responding to membrane stress.

Further research into these newly described systems will yield broadly applicable mechanistic insights, given their conservation across life. For example, SHOCT domains are understudied since they are thought to be absent in model organisms (96). However, our approach identified divergent homologs in the model organism *B. subtilis* (<u>CAB15517.1</u>) that may prompt new studies. The extensive findings from applying gene neighborhood and domain architecture searches support their viability, and such methods help reduce the space of annotated hypothetical proteins without classified domains and add functional clues. Finally, we provide another pillar of support to recent conclusions that PspA/Snf7 homologs were already present in the last universal common ancestor (19, 20), and deepen our understanding of the origins and diversification of all life. All the findings (data, results, and visual summaries) from this work are found in an interactive and queryable companion web application available at https://jravilab.org/psp.

## Methods

### Query and subject selection

We queried classical PSP members — PspA (eight representatives across Proteobacteria, Actinobacteria, Firmicutes (two paralogs), Cyanobacteria, and Archaea); PspM (Rv2743c) and PspN (Rv2742c) (from *M. tuberculosis*); PspB and PspC (from *E. coli*); LiaI, LiaG, and LiaF (from *B. subtilis*) — against all sequenced genomes across the tree of life. Transcription factors and two-component systems, including PspF (from *E. coli*), ClgR (from *M. tuberculosis*), LiaRS (from *B. subtilis*), are not part of this detailed analyses or the web app, due to their prevalence across the bacterial kingdom. We ran homology searches for the other representative PSP components against custom databases of ∼6500 completed representative or reference genomes with taxonomic lineages or the NCBI non-redundant database (**Tables S1** and **S2**) (108, 109). All PSP homologs are listed in the <u>web app</u>, with representatives in **Tables S3** and **S4**. We obtained the phyletic order (sequence) from NCBI taxonomy and BV-BRC (108–112).

### Identification and characterization of protein homologs

We analyzed the classical PSP proteins from the best-studied operons *pspF||ABCDE-G* (from *E. coli*), *liaIHGFSR* (from *B. subtilis*), and *clgRpspAMN* (from *M. tuberculosis*) and their phyletic spread to locate and stratify all PSP homologs. We first resolved these PSP proteins into their constituent domains and used each individual domain as a query for all our phylogenetic analyses to ensure an exhaustive search and identification of all related PSP proteins, including remote homologs. This approach allows us to find homologies beyond those from full-length proteins only. We then performed homology searches for each constituent domain across the tree of life (approximately 6500 representative or reference genomes). We used a combination of protein domain and orthology databases, iterative homology searches, and multiple sequence alignments to detect domains, signal peptides, and transmembrane (TM) regions to help construct the domain architecture of each query PSP protein. The ubiquitous presence of transcription factor HTH domains and histidine kinase-receiver domain two-component systems in bacterial phyla precluded dedicated searches with these domains; instead, we noted their occurrence in predicted operons alongside core PSP genes to identify transcriptional regulation partners.

We ensured the identification of a comprehensive set of homologs (close and remote) for each queried protein via iterative searches using PSI-BLAST (113) with sequences of both full-length proteins and the corresponding constituent domains. More distant relationships were determined using profile-profile searches with the HHpred program (114). Protein searches were conducted using homologous copies from multiple species as starting points. We aggregated search results and recorded the numbers of homologs per species and genomes carrying each of the query proteins (<u>web app</u>). These proteins were clustered into orthologous families using the similarity-based clustering program BLASTCLUST (115). SignalP, TMHMM, Phobius, JPred, Pfam, and custom profile databases (116–121) were used to identify signal peptides, TM regions, known domains, and the secondary protein structures in every genome. Homolog information, including domain architectures, is available in the <u>web app</u> (‘Data,’ ‘Domain architectures’ tabs).

### Neighborhood search

Bacterial gene neighborhoods (±7 genes flanking each protein homolog) were retrieved using a Perl script from GenBank (108, 109). Gene orientation, domains, and secondary structures of the neighboring proteins were characterized using the same methods applied to query homologs above. Genomic contexts are available in the <u>web app</u> (‘Genomic contexts’ tab). We note that eukaryotic components (*e.g.,* Snf7) appearing alone are an artifact of the genomic context analysis being restricted (and relevant) to bacteria and archaea (see <u>web app</u>).

### Phylogenetic analysis

Multiple sequence alignment (MSA) of the identified homologs was performed using Kalign (122) and MUSCLE (123) with default parameters. Domain/gene trees were constructed with FastTree 2.1 with default parameters (124) and visualized and labels colored in FigTree v1.4.4 (125). In parallel, the PspA/Snf7 MSA was also passed to ModelTest-NG, which reported the best performing substitution model (LG+I+G4+F) (126, 127), and then RAxML-NG-MPI (v1.2.0) (128) on CU’s Alpine HPC cluster and a maximum likelihood phylogenetic tree constructed (--all, –model LG+I+G4+F, –tree rand{25} pars{25}, –bs-trees autoMRE, –threads 60). Bootstrapping was performed until RAxML-NG assessed statistical convergence at 480 replicates (**Fig. S2**). Phylogenetic trees are available in the <u>web app</u> (‘Phylogeny’ tab).

### Network reconstruction

The PSP proximal neighborhood network was constructed on the domain architectures and genomic contexts of PspA and its cognate partner proteins (**Tables S1** and **S2**). The nodes represented the domains, and edges indicated a shared neighborhood (domains of the same protein or neighboring proteins). Proximity networks are available in the <u>web app</u> (‘Domain architectures’ tab).

### Web application

We built our interactive and searchable web application using R Shiny (129). All data analyses and visualizations were carried out using R or RStudio (130, 131) with R packages (132–144). Data and results from our study are available on the web application that serves as a companion data summarization and visualization dashboard to this manuscript (https://jravilab.org/psp). The web app can be queried with the protein accession numbers, domain names, or species/lineages cited inline throughout the Results section. While not used directly in this analysis, our MolEvolvR protein analysis web app can be used with the accession numbers of any protein we describe for readers who would like to further explore a protein’s domain architectures and phylogenetic distribution across life (https://jravilab.org/molevolvr) (104).

## Declarations

## Acknowledgments

We are incredibly grateful to Arjun Krishnan and Krishnan Raghunathan for detailed comments on the manuscript and web application, guidance, and support; Emily Meyer and Jerome McKay for critical edits of the manuscript. We also thank Srinand Sreevatsan, Beronda Montgomery, Deborah Johnson, and the MSU Diversity Research Network for support and writing spaces. Finally, we thank members of the JRaviLab for feedback on the web application, and especially Jacob Krol and Faisal Alquaddoomi for their additional help with the final deployment. This work utilized the Alpine high-performance computing resource at the University of Colorado Boulder. Alpine is jointly funded by the University of Colorado Boulder, the University of Colorado Anschutz, Colorado State University, and the National Science Foundation (award: 2201538).

## Author Contributions

J.R. and M.L.G. conceived the study; J.R., V.A., L.A., and M.L.G. designed the study; J.R. and V.A acquired the data, performed all the analyses, and made the figures and tables. J.R., V.A., L.A., and M.L.G. interpreted the results and wrote the manuscript. S.Z.C built the web application with all the results (data summarization and visualization) with J.R.; S.Z.C. also contributed to making figures and linking identifiers in the manuscript to reference databases. E.P.B. contributed to writing the manuscript, performed phylogenetic tree construction, and mapped Pfam to Interpro domain IDs throughout the manuscript. P.D. contributed to renaming some domains.

### Data Availability and Reuse

All the data, analyses, and visualizations are available in our interactive and queryable web application: https://jravilab.org/psp. Text, figures, and the <u>web app</u> are licensed under Creative Commons Attribution CC BY 4.0.

