## Supplementary Material for "The Phage-shock-protein (PSP) Envelope Stress Response: Discovery of Novel Partners and Evolutionary History"

#### Table of Contents

**Supplementary Figures and Tables**

**Supplementary Figures**

Figure S1. Mutual exclusion phyletic pattern of PspM, Thioredoxin

Figure S2. PspA maximum likelihood gene tree with bootstrap values by RAxML-NG.

**Supplementary Tables**

Table S1. Summary of PSP partner domains (query proteins)

Table S2. Summary of novel PSP partner domains

Table S3. Domain architectures, genomic contexts, and lineages of representative PspA/Snf7 homologs. (next pages)

Table S4. Domain architectures, genomic contexts, and lineages of representative homologs of PSP cognate partner domains. (next pages)

**Supplementary Text**

1. Domain definitions

1.1. LiaI-LiaF-TM and Toastrack domains

1.2 PspM and PspN domains

1.3 PspAA and PspAB domains

2. Novel PSP associations

2.1 PspA/Snf7 domain architectures

2.2 Vps4 and $AAA^{+}$-ATPases

2.3 PspA with PspM or Thioredoxin

2.4 Novel contexts containing Toastrack

Toastrack and TM domains

Toastrack and transcription factors

**References**

### Supplementary Figures and Tables

*All PSP results (data summarizations and visualizations) can be accessed via our easy-to-use interactive web app: [https://jravilab.org/psp](https://jravilab.org/psp).*

#### Supplementary Figures

##### Figure S1. Mutual exclusion phyletic pattern of PspM, Thioredoxin

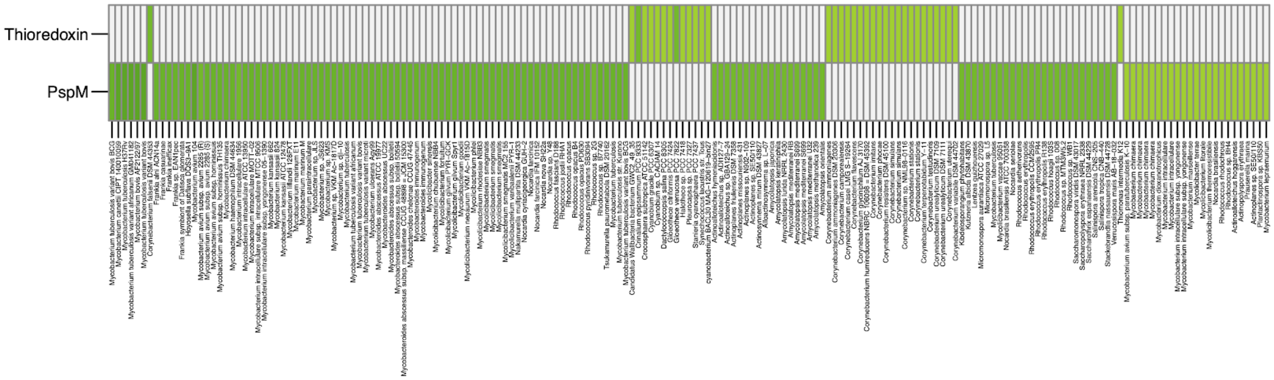

**Figure S2. PspA maximum likelihood gene tree with bootstrap values by RAxML-NG.**

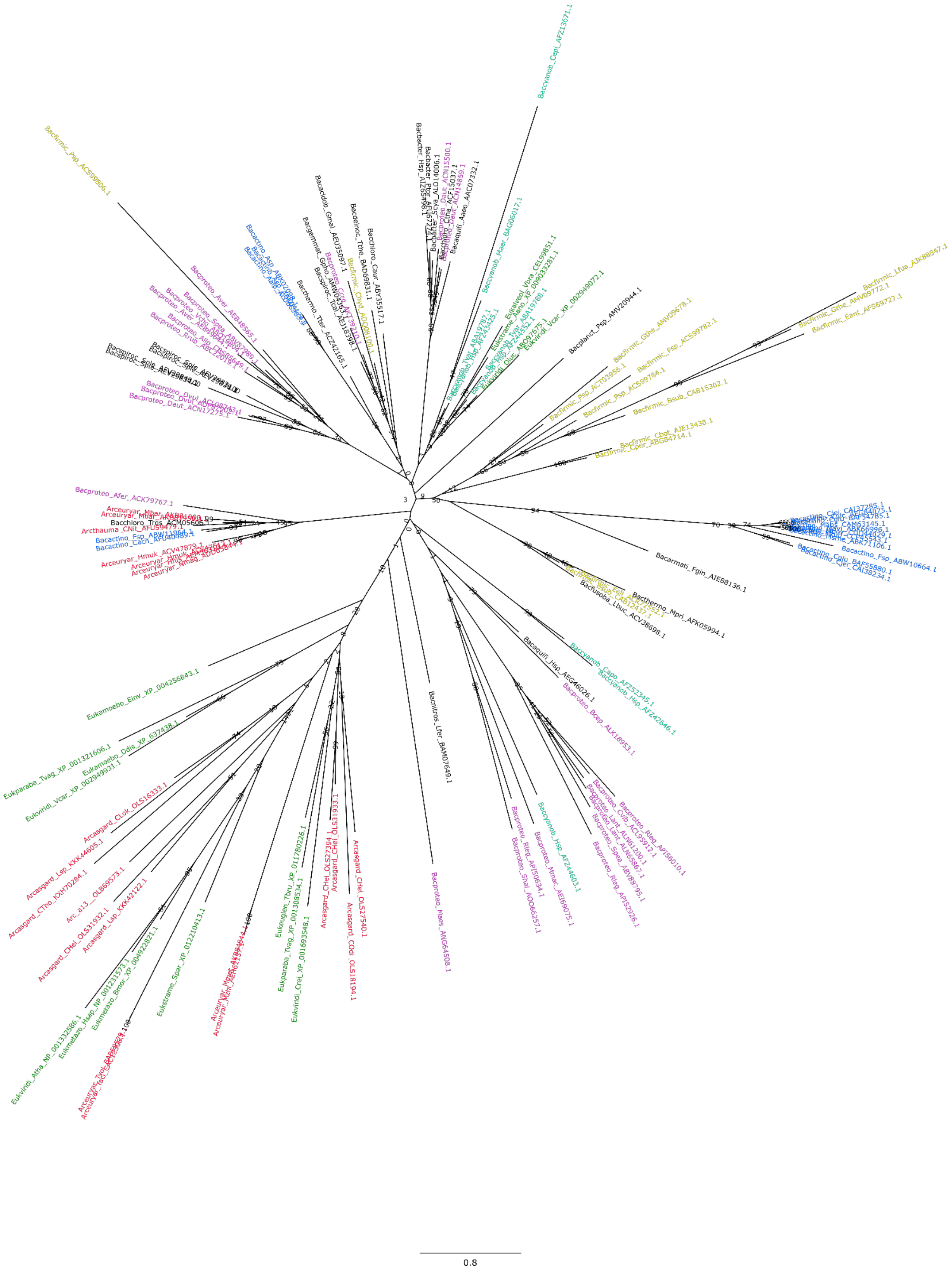

#### Supplementary Tables

**Table S1. Summary of PSP partner domains (query proteins)**

| Domain Name | Old Name/Aliases | Pfam | PDB |
|---|---|---|---|
| PspA | PspA_IM30<br>LiaH, Vipp1, YjfJ, YdjF | PF04012 | 4WHE |
| Snf7 | | PF003357 | 5FD7 |
| PspA/Snf7 | PspA/ESCRT-III | CL0235 | 3FRV |
| Toastrack | DUF4097/DUF2154/DUF2807<br>LiaF/LiaG, YvlB, YthC, Toastrack_N (N-terminal region of Toastrack; PF17115) | PF13349/<br>PF09922/<br>PF10988/PF17115 | 4QRK/<br>4OPW |
| LiaI-LiaF-TM | DUF2157<br>LiaI/LiaF | PF09925 | |
| PspM | Rv2743c | | |
| PspN | Rv2742c | | |
| PspN_N | N-terminal region of PspN | | |
| DUF3046 | C-terminal region of PspN | PF11248 | |
| PspB | | PF06667 | |
| PspC | YthA/YthB, YvlC | PF04024 | |

**Table S2. Summary of novel PSP partner domains**

| Domain Name | Old Name/Aliases | Pfam | PDB |
|---|---|---|---|
| HAAS | DUF1700 alpha-helical/<br>DUF1129/Yip1/<br>DUF1048 | PF08006/PF06570/<br>PF04893/PF06304 | 2O3L |
| SHOCT-bihelical | DUF1707, SHOCT | PF08044/PF09851 | |
| PspAA | PspA-associated domain A | | |
| PspAB | PspA-associated domain B | | |
| Vps4-AAA-ATPase/<br>Classical-AAA-ATPase | | PF08432/<br>PF00004 | 5FVK/<br>3U5Z |
| MIT | | PF04212 | 5FVK |

| | | | |
|---|---|---|---|
| Thioredoxin | | PF00085 | 2OE3 |
| ClgR-HTH | XRE-HTH/HTH_3, ClgR, cHTH | PF01381 | 6IRP |
| TM-Flotillin dyad | Flot | PF15975 | |
| Band-7 | Band_7 | PF01145 | 3BK6 |
| Spermine synthase | Spermine_synth | PF01564 | 6O63 |
| YjfL-TM(s) | DUF350 | PF03994 | |
| CesT_Tir | CesT | PF05932 | 1TTW |
| CesT_Tir-DUF2170 | DUF2170 | PF09938 | |
| TPM_phosphatase | | PF04536 | 4OA3 |
| SHOCT-like | DUF1707 | PF08044 | |
| Caspase | Peptidase_C14 | PF00656 | 3UO8 |
| PadR-like_wHTH PadR-wHTH | PadR-HTH/PadR | PF03551 | 1XMA |
| RHH | TraJ-RHH/RHH_1 | PF01402 | 3OD2 |
| SIGMA-HTH | | | |
| GerE-HTH/ DUF2089-HTH | GerE/ DUF2089 | PF00196/PF09862 | 2JPC |
| GNTR-HTH | GntR | PF00392 | 4R1H |

**Table S3. Domain architectures, genomic contexts, and lineages of representative PspA/Snf7 homologs. (next pages)**

**Table S4. Domain architectures, genomic contexts, and lineages of representative homologs of PSP cognate partner domains. (next pages)**

<table>
<tr><th colspan="5">Table S3: Representative PspA/Snf7 homologs<br>Gene, Lineage information and Genomic Contexts grouped by Domain Architectures</th></tr>
<tr><th colspan="2">Gene Info</th><th colspan="2">Lineage Info</th><th></th></tr>
<tr><th>AccNum</th><th>GeneName</th><th>Species</th><th>Lineage</th><th>GenContext</th></tr>
<tr><td colspan="5">MIT+Vps4-AAA-ATPase</td></tr>
<tr><td>CKH37208.1</td><td>ftsH_1</td><td>Mycolicibacterium smegmatis</td><td>bacteria>actinobacteria</td><td>MIT+Vps4-AAA-ATPase-></td></tr>
<tr><td>ACB74714.1</td><td>Oter_1429</td><td>Opitutus terrae PB901</td><td>bacteria>PVC_group>verrucomicrobia</td><td>MIT+Vps4-AAA-ATPase->TPR+TM(s)-></td></tr>
<tr><td colspan="5">NlpC+PspA</td></tr>
<tr><td>AFZ52345.1</td><td>Cyan10605_0189</td><td>Cyanobacterium aponinum PCC 10605</td><td>bacteria>cyanobacteria</td><td>NlpC+PspA-></td></tr>
<tr><td colspan="5">PspA</td></tr>
<tr><td>AKJ06548.1</td><td>AA314_08174</td><td>Archangium gephyra</td><td>bacteria>proteobacteria>deltaproteobacteria</td><td><-ABC-ATPase||Glycos_trans_3N+Glycos_transf_3+PYNP_C->X(s)->PspA->SIG+PBPB->SIG+TM+Snf7->||<-X||SIG+DUF3352->ABC-ATPase->SIG+TM(s)->SIG+TM(s)-></td></tr>
<tr><td>AEY64321.1</td><td>Clo1100_0028</td><td>Clostridium sp BNL1100</td><td>bacteria>firmicutes</td><td><-ParA-Soj-PloopNTPase||X(s)->GNTR-HTH->inactive-Classical-AAA+Classical-AAA->PspA->ACET-></td></tr>
<tr><td>ANQ40502.1</td><td>BAR24_02900</td><td>Gluconobacter oxydans</td><td>bacteria>proteobacteria>alphaproteobacteria</td><td><-PspF-NtrC-AAA+FIS-HTH||PspA->PspB->TM+Toastrack->TM+Toastrack->SIG+TM(s)->ABC-ATPase-></td></tr>
<tr><td>AOL22920.1</td><td>Ga0102493_111899</td><td>Erythrobacter litoralis</td><td>bacteria>proteobacteria>alphaproteobacteria</td><td><-PspF-NtrC-AAA+FIS-HTH||X->PspA->PspB->PspC+PspB->SIG+PspB->SIG+PspB->SIG+PspB->SIG+PspB->PspB-></td></tr>
<tr><td>BAB38581.1</td><td>yjfJ</td><td>Escherichia coli O157:H7 str Sakai</td><td>bacteria>proteobacteria>gammaproteobacteria</td><td>CesT_Tir-DUF2170->PspA->DUF2491->YjfL-TM+TM(s)->LipoSIG+DUF1190->SpermGS-ATPgrasp-></td></tr>
<tr><td>AMJ95269.1</td><td>AVL56_13815</td><td>Alteromonas addita</td><td>bacteria>proteobacteria>gammaproteobacteria</td><td>CesT_Tir-DUF2170->PspA->Ion_trans_2+TrkA_N+TrkA_C->DUF2491->YjfL-TM+TM(s)->LipoSIG+DUF1190->SpermGS-ATPgrasp-></td></tr>
<tr><td>ABK71106.1</td><td>MSMEG_2695</td><td>Mycolicibacterium smegmatis MC2 155</td><td>bacteria>actinobacteria</td><td>ClgR-HTH->PspA->PspM->||<-X(s)->DUF3046-></td></tr>
<tr><td>CCP45543.1</td><td>35kd_ag</td><td>Mycobacterium tuberculosis H37Rv</td><td>bacteria>actinobacteria</td><td>ClgR-HTH->PspA->PspM->PspN_N+DUF3046->||<-X(s)->DUF3046-></td></tr>
<tr><td>AOS62694.1</td><td>TL08_09395</td><td>Actinoalloteichus hymeniacidonis</td><td>bacteria>actinobacteria</td><td>ClgR-HTH->UA74_09550-lowcomplexity->PspA->PspM-></td></tr>
<tr><td>ANX06812.1</td><td>AS891_06225</td><td>Bacillus subtilis subsp subtilis</td><td>bacteria>firmicutes</td><td>LiaI-LiaF-TM->PspA->TM+Toastrack->LiaI-LiaF-TM+Toastrack->SIG+TM+HAMP+HISKIN->REC+wHTH-></td></tr>
<tr><td>AAN56746.1</td><td>SO_3765</td><td>Shewanella oneidensis MR1</td><td>bacteria>proteobacteria>gammaproteobacteria</td><td>LipoSIG+Ctha_1186+Low-comp->YjfL-TM+TM(s)->SIG+TM(s)+Spermine_synth->CesT_Tir-DUF2170->PspA->DUF4178->RHH-></td></tr>
<tr><td>AFZ14666.1</td><td>Cri9333_3857</td><td>Crinalium epipsammum PCC 9333</td><td>bacteria>cyanobacteria</td><td>PspA->PspA->Thioredoxin-></td></tr>
<tr><td>AAM04874.1</td><td>MA_1460</td><td>Methanosarcina acetivorans C2A</td><td>archaea>euryarchaeota</td><td>PspA->PspAA-></td></tr>
<tr><td>CAB51252.1</td><td>SCO2168</td><td>Streptomyces coelicolor A32</td><td>bacteria>actinobacteria</td><td>PspA->PspAA->SIG+TM(s)+HISKIN->REC+wHTH-></td></tr>
<tr><td>ABW11964.1</td><td>Franean1_2534</td><td>Frankia sp EAN1pec</td><td>bacteria>actinobacteria</td><td>PspA->PspAA->TM(s)+Metallopeptidase+TM(s)->PspAB-></td></tr>
<tr><td>AEN65073.1</td><td>Entas_2342</td><td>Enterobacter soli</td><td>bacteria>proteobacteria>gammaproteobacteria</td><td>PspA->PspB->PspC+PspB->PspD->DO-GTPase2->TM(s)+IIGP1->PspF-NtrC-AAA+FIS-HTH-></td></tr>
<tr><td>ANW39986.1</td><td>A9L45_10880</td><td>Escherichia coli O157:H7</td><td>bacteria>proteobacteria>gammaproteobacteria</td><td>PspA->PspB->PspC+PspB->PspD->PspE-SIG+RHOD-CDC25-></td></tr>
<tr><td>ANX09535.1</td><td>AS891_20665</td><td>Bacillus subtilis subsp subtilis</td><td>bacteria>firmicutes</td><td>PspA->ZnR(s)+TM->SIG+TPM_phosphatase+TM->Band-7+ZnR-></td></tr>
<tr><td>CBH24266.1</td><td>SRM_01345</td><td>Salinibacter ruber M8</td><td>bacteria>FCB_group>bacteroidetes</td><td>Ribosomal_L31->Glycos_trans_3N+Glycos_transf_3+PYNP_C->PspA->Rmar_0091-Coiled-coil->SIG+TM+Snf7->||<-X<-MoaC</td></tr>
<tr><td>AKX93460.1</td><td>MOTHE_c06560</td><td>Moorella thermoacetica</td><td>bacteria>firmicutes</td><td>SIG+SHOCT-bihelical->TM(s)+Metallopeptidase+TM(s)->PspAB->PspA->PspAA-></td></tr>
<tr><td>APB74393.1</td><td>PPYC2_05025</td><td>Paenibacillus polymyxa</td><td>bacteria>firmicutes</td><td>SIG+TM(s)->LiaI-LiaF-TM->PspA->PspC+Coiled-coil->PspA->SIG+TM+Toastrack->SIG+TM+HISKIN->REC+wHTH-></td></tr>
<tr><td>AKK09942.1</td><td>CTEST_12695</td><td>Corynebacterium testudinoris</td><td>bacteria>actinobacteria</td><td>Thioredoxin->PspA-></td></tr>
<tr><td>ANH61663.1</td><td>I597_2772</td><td>Dokdonia donghaensis DSW1</td><td>bacteria>FCB_group>bacteroidetes</td><td>TM(s)->PadR-like-wHTH->HAAS+PspC+LiaI-LiaF-TM+Toastrack->SIG+NTF2->TM+Toastrack->||<-CHTH+Protease||CesT_Tir->PspA->YqiJ-YuaF-SIG+TM(s)->SIG+Band-7+Coiled-coil+TM-Flotillin->Betapropeller+Coiled-coil+AAA-ATPase-></td></tr>
<tr><td>AKB54760.1</td><td>MSBRM_1762</td><td>Methanosarcina barkeri MS</td><td>archaea>euryarchaeota</td><td>TM(s)+Metallopeptidase+TM(s)->PspAB->||<-X(s)->PspA->PspAA-></td></tr>
<tr><td colspan="5">PspA(s)</td></tr>
<tr><td>BAG06017.1</td><td>MAE_61950</td><td>Microcystis aeruginosa NIES843</td><td>bacteria>cyanobacteria</td><td>PspA->PspA->PspA(s)-></td></tr>
<tr><td colspan="5">PspA+PspAA</td></tr>
<tr><td>ACU53894.1</td><td>Afer_0955</td><td>Acidimicrobium ferrooxidans DSM 10331</td><td>bacteria>actinobacteria</td><td>PspA+PspAA-></td></tr>
<tr><td colspan="5">PspAB</td></tr>
<tr><td>AAZ55047.1</td><td>Tfu_1009</td><td>Thermobifida fusca YX</td><td>bacteria>actinobacteria</td><td><-TIMbarrel<-X||PspA->PspAA->TM(s)+Metallopeptidase+TM(s)->PspAB-></td></tr>
<tr><td colspan="5">SIG+MMPL+Snf7+MMPL</td></tr>
<tr><td>CAM62382.1</td><td>MAB_2301</td><td>Mycobacteroides abscessus ATCC 19977</td><td>bacteria>actinobacteria>actinobacteria</td><td>Mycobact_memb->SIG+MMPL+Snf7+MMPL-></td></tr>
<tr><td colspan="5">SIG+TM+Snf7</td></tr>
<tr><td>OGG56892.1</td><td>A3F84_10925</td><td>Candidatus Handelsmanbacteria bacterium RIFCSPLOWO2_12_FULL_64_10</td><td>bacteria</td><td>FAD_binding_5+CO_deh_flav_C->X(s)->PspA->SIG+PBPB+OmpA->SIG+TM(s)->ABC-ATPase->SIG+TM+Snf7->inactive-Classical-AAA+Classical-AAA-></td></tr>
<tr><td colspan="5">Snf7</td></tr>
<tr><td>OLS27540.1</td><td>HeimC3_03190</td><td>Candidatus Heimdallarchaeota archaeon LC_3</td><td>archaea>asgard_group</td><td>MIT+Vps4-AAA-ATPase->||<-X(s)->Snf7->Snf7->MIT+Vps4-AAA-ATPase->ESCRT-II-></td></tr>
<tr><td>CBY21170.1</td><td>GSOID_T00008924001</td><td>Oikopleura dioica</td><td>eukaryota>metazoa>chordata</td><td>Snf7->Snf7->Snf7-></td></tr>
</table>

More information available on our webapp.

Table S4: Representative homologs of Psp cognate partner domains
Gene, Lineage information and Genomic Contexts grouped by Domain Architectures

| Gene Info | | Lineage Info | | |
|---|---|---|---|---|
| AccNum | GeneName | Species | Lineage | GenContext |
| **DUF4178** | | | | |
| AAN56747.1 | SO_3766 | Shewanella oneidensis MR1 | bacteria>proteobacteria>gammaproteobacteria | LipoSIG+Ctha_1186+Low-comp->YjfL-TM+TM(s)->SIG+TM(s)+Spermine_synth->CesT_Tir-DUF2170->PspA->DUF4178->RHH-> |
| **HAAS+FTSW_RODA_SPOVE** | | | | |
| CAC98500.1 | lmo0421 | Listeria monocytogenes EGDe | bacteria>firmicutes | SIGMA-Factor->PadR-like-wHTH->HAAS+FTSW_RODA_SPOVE-> |
| **HAAS+MacB_PCD+FtsX+MacB_PCD+FtsX** | | | | |
| ACO32024.1 | ACP_2125 | Acidobacterium capsulatum ATCC 51196 | bacteria>acidobacteria | PadR-like-wHTH->HAAS+MacB_PCD+FtsX+MacB_PCD+FtsX-> |
| **HAAS+Pentapeptide** | | | | |
| AOH56696.1 | ABE28_020195 | Bacillus muralis | bacteria>firmicutes | PadR-like-wHTH->HAAS+Pentapeptide-> |
| **LiaI-LiaF-TM** | | | | |
| APB74392.1 | PPYC2_05020 | Paenibacillus polymyxa | bacteria>firmicutes | SIG+TM(s)->LiaI-LiaF-TM->PspA->PspC+Coiled-coil->PspA->SIG+TM+Toastrack->SIG+TM+HISKIN->REC+wHTH-> |
| **LiaI-LiaF-TM+Toastrack** | | | | |
| ABD83157.1 | Sde_3902 | Saccharophagus degradans 240 | bacteria>proteobacteria>gammaproteobacteria | LiaI-LiaF-TM+Toastrack->SIG+TM(s)+HISKIN->REC+wHTH-> |
| AFH48155.1 | IALB_0443 | Ignavibacterium album JCM 16511 | bacteria>ignavibacteriae | LiaI-LiaF-TM+Toastrack->SIG+TM(s)+HISKIN->REC+wHTH->X(s)->TM-> |
| AGK93623.1 | LA14_0400 | Lactobacillus acidophilus La14 | bacteria>firmicutes | LytTR->LiaI-LiaF-TM+Toastrack-> |
| CAL82154.1 | CBO0601 | Clostridium botulinum A str ATCC 3502 | bacteria>firmicutes | LytTR->LiaI-LiaF-TM+Toastrack-> |
| **PspC** | | | | |
| CAB15516.1 | yvlC | Bacillus subtilis subsp subtilis str 168 | bacteria>firmicutes | yvlA-SIG+TM(s)->SHOCT-like+Toastrack->PspC->SIG+TM(s)->‖<-SIG+TM(s) |
| **PspC+LiaI-LiaF-TM** | | | | |
| ABY34522.1 | Caur_1294 | Chloroflexus aurantiacus J10fl | bacteria>chloroflexi | PspC+LiaI-LiaF-TM-> |
| **PspC+LiaI-LiaF-TM+HISKIN** | | | | |
| AIJ13865.1 | SLIV_14405 | Streptomyces lividans TK24 | bacteria>actinobacteria | <-PspC+TM(s)+Toastrack‖PspC+LiaI-LiaF-TM+HISKIN->REC+wHTH-> |
| **PspC+LiaI-LiaF-TM+TM(s)+Toastrack** | | | | |
| ACV77657.1 | Namu_1253 | Nakamurella multipartita DSM 44233 | bacteria>actinobacteria | <-REC+wHTH<-PspC+LiaI-LiaF-TM+HISKIN‖PspC+LiaI-LiaF-TM+TM(s)+Toastrack->X->‖<-SIG+TM(s) |
| **PspC+TM(s)+Toastrack** | | | | |
| AIJ13866.1 | SLIV_14410 | Streptomyces lividans TK24 | bacteria>actinobacteria | <-REC+wHTH<-PspC+LiaI-LiaF-TM+HISKIN‖PspC+TM(s)+Toastrack-> |
| **SHOCT-bihelical+Toastrack** | | | | |
| CCP43715.1 | Rv0966c | Mycobacterium tuberculosis H37Rv | bacteria>actinobacteria | SHOCT-bihelical+Toastrack-> |
| CAB88834.1 | SCO2893 | Streptomyces coelicolor A32 | bacteria>actinobacteria | TM(s)->TM(s)->ABC-ATPase->SHOCT-bihelical+Toastrack-> |
| **SHOCT-like+Toastrack** | | | | |
| CAB15517.1 | yvlB | Bacillus subtilis subsp subtilis str 168 | bacteria>firmicutes | yvlA-SIG+TM(s)->SHOCT-like+Toastrack->PspC->SIG+TM(s)->‖<-SIG+TM(s) |
| **SIG+LiaI-LiaF-TM+TM+Toastrack** | | | | |
| AEU34960.1 | AciX8_0610 | Granulicella mallensis MP5ACTX8 | bacteria>acidobacteria | SIGMA-Factor->anti-sigma-ZF+TM->TM+LiaI-LiaF-TM->SIG+LiaI-LiaF-TM+TM+Toastrack-> |
| **SIG+LiaI-LiaF-TM+Toastrack** | | | | |
| CAB15300.1 | liaF | Bacillus subtilis subsp subtilis str 168 | bacteria>firmicutes | LiaI-LiaF-TM->PspA->TM+Toastrack->LiaI-LiaF-TM+Toastrack->SIG+TM+HAMP+HISKIN->REC+wHTH-> |
| **SIG+TM(s)+Toastrack** | | | | |
| ABD31150.1 | SAOUHSC_02100 | Staphylococcus aureus subsp aureus NCTC 8325 | bacteria>firmicutes | SIG+TM(s)+Toastrack->SIG+TM(s)+HISKIN->REC+wHTH->‖<-SIG+TM(s) |
| **TM+DUF4178** | | | | |
| CCP45393.1 | Rv2597 | Mycobacterium tuberculosis H37Rv | bacteria>actinobacteria | TM+DUF4178->DUF2617->SIG+DUF4247->YjfL-TM+TM(s)->SIG+TM(s)+Spermine_synth-> |
| **TM+Toastrack** | | | | |
| AJI31452.1 | BF28_3762 | Bacillus cereus E33L | bacteria>firmicutes | ABC-ATPase->TM(s)->TM+Toastrack-> |
| AAM36414.1 | XAC1545 | Xanthomonas citri pv citri str 306 | bacteria>proteobacteria>gammaproteobacteria | GNTR-HTH->ABC-ATPase->TM(s)->TM+Toastrack->SIG+DUF2884-> |
| AFK03672.1 | Emtol_2536 | Emticicia oligotrophica DSM 17448 | bacteria>FCB_group>bacteroidetes | SIGMA-Factor->anti-sigma-ZF+TM+HEAT->TM+Toastrack->TM+Toastrack-> |
| **TM+Toastrack+CASPASE** | | | | |
| AFY83227.1 | Oscil6304_3666 | Oscillatoria acuminata PCC 6304 | bacteria>cyanobacteria | TM+Toastrack+CASPASE-> |
| **ZnR+DUF2089-HTH+SHOCT-like** | | | | |
| ADE70705.1 | BMQ_3692 | Bacillus megaterium QM B1551 | bacteria>firmicutes | ZnR+DUF2089-HTH+SHOCT-like->SHOCT-like+X-> |
| **ZnR+PspC** | | | | |
| ABC83427.1 | Adeh_3661 | Anaeromyxobacter dehalogenans 2CPC | bacteria>proteobacteria>deltaproteobacteria | ZnR+PspC-> |

More information available on our webapp.

### Supplementary Text

#### 1. Domain definitions

##### 1.1. LiaI-LiaF-TM and Toastrack domains

To best characterize the LiaIGF proteins, we used PSI-BLAST searches from the three sub-sequences of the full-length proteins, N-terminal TM region of LiaF, the C-terminus globular domain (DUF2154) of LiaF, and the globular domain in LiaG (DUF4097), followed by structure-informed sequence alignment. These analyses revealed that LiaI and LiaG bear remarkable similarities to the N-terminal TM and C-terminal globular regions of the LiaF protein, respectively. We discovered that the globular domains of these LiaG–LiaF proteins are homologs of each other and that the profiles detected by Pfam in this region, DUF2154, DUF4097, and DUF2807, can be unified into a single domain called "**Toastrack**" that has a single-stranded right-handed beta-helix fold with a unique N-terminal 7-stranded region that displays a complex intertwining of strands (PDB: 4QRK; Toastrack_N, Pfam: PF17115 only has this unique N-terminal region) [**Fig. 1A**; **Table S1**]. Likewise, the homology between the 4TM (four TM) regions of LiaI and the N-terminal domain DUF2157 of LiaF led us to rename the 4TM region as '**LiaI-LiaF-TM**' [**Fig. 1B**; **Table S1**]. Thus, the results of our analyses define two new domains: LiaI-LiaF-TM and Toastrack.

##### 1.2 PspM and PspN domains

PspM comprises two TM regions and no other distinct domains. PspN contains a short domain at the C-terminus, DUF3046, and a yet uncharacterized N-terminal domain, which we now call **PspN_N**. We found that DUF3046 is α-helical with highly conserved threonine and cysteine residues that might be required for its function.

To further characterize the DUF3046 homologs, we used nucleotide sequences rather than translated open reading frames (ORFs), followed by sequence alignment analysis [**Fig. 1B**]. We found that the DUF3046 domain, which is widespread across Actinobacteria, is more similar to the short downstream protein, Rv2738c, than to the C-terminus of the fourth member of the *M. tuberculosis* operon, PspN (encoded by *Rv2742c*).

##### 1.3 PspAA and PspAB domains

The PspA neighborhood analysis identified a new component in the proximity of PspA, which is a protein containing a novel trihelical domain (with absolutely conserved R and D) present in Euryarchaeota, Thaumarchaeota, Actinobacteria, Chloroflexi, Firmicutes, and a few Alpha- and Gammaproteobacteria). This protein occurs in a two-gene cluster with PspA [**Fig. 3**; **Table S3**; *e.g.,* MA_1460; AAM04874.1, Methanosarcina]. This domain mostly occurs by itself but is occasionally fused to an N-terminal PspA in Actinobacteria and Chloroflexi [**Fig. 3**; **Table S3**; *e.g.,* ACU53894.1, Acidimicrobium]. We call this domain **PspAA** (for **PspA-associated domain A**; **Table S2**; web app). In a few bacterial and archaeal lineages, the PspA–PspAA dyad co-occurs with another dyad comprising a membrane-associated Metallopeptidase and a protein with a novel domain, which we termed **PspAB** (for **PspA-associated domain B**, AAZ55047.1, Tfu_1009 Thermobifida) [**Fig. 3**; **Table**

**S2**; web app]. This predicted operon occasionally contains a third gene coding for a SHOCT-like bihelical domain-containing protein in various bacterial and archaeal lineages [**Fig. 3**; **Table S3**; ABW11964.1, *Frankia*; AKB54760.1, *Methanosarcina*; AKX93460.1, *Moorella*].

#### 2. Novel PSP associations

##### 2.1 PspA/Snf7 domain architectures

A very small fraction of PspA homologs shows variation in their domain architecture (proteins that contain fusions with PspA instead of carrying PspA alone). For example, cyanobacterial PspA homologs show some interesting variations: a few have dyads or triads of PspA, either as repeated domains within a polypeptide or a predicted operon with multiple copies of PspA-containing genes [**Fig. 3**; *e.g.,* BAG06017.1], while others carry an additional hydrolase domain of NlpC/P60 superfamily at the N-terminus that is predicted to catalyze the modification of phosphatidylcholine, thus altering membrane composition [**Fig. 3**; **Table S3**; AFZ52345.1;(1)]. We also find a novel fusion of PspA with **PspAA** in Actinobacteria (ACU53894.1, *Acidimicrobium*; defined in the section on PspAA below). Similar to the PspA homologs, a search for the related superfamily, Snf7, revealed minimal variation in domain architecture, with occasional fusions (<5%) found only in eukaryotes [**Fig. 2B**]. Some Actinobacteria, such as *Mycobacteroides abscessus*, have an Snf7 homolog [CAM62382.1, **Fig. 2B**] fused to an RND-family transporter member. The latter transports lipids and fatty acid and is flanked by two genes encoding the *Mycobacterium*-specific TM protein with a C-terminal Cysteine-rich domain (2).

##### 2.2 Vps4 and AAA$^+$-ATPases

One or more copies of an *snf7* gene [*e.g.,* OLS27540.1; **Table S3**] and a gene for the **VPS4-like AAA$^+$-ATPase** (with an N-terminal MIT domain and C-terminal oligomerization domain; **Table S2**) are known to occur together in Archaea; they define the core of an ESCRT complex (3). However, we observed some diversity between different archaeal lineages. For example, the Asgardarchaeota contain a genomic context that is most similar to eukaryotes. This archaeal context is composed of the Vps4 AAA$^+$-ATPase and Snf7-encoding genes along with an ESCRT-II gene that encodes a protein with multiple winged helix-turn-helix (wHTH) domains (4). In Crenarchaeota, Snf7 and the Vps4 AAA+-ATPase are encoded in a distinct three-gene operon, which contains a gene coding for a CdvA-like coiled-coil protein with an N-terminal PRC-barrel domain implicated in archaeal cell division (5). In this case, the Snf7 domain is fused to a C-terminal wHTH domain, which might play a role equivalent to the ESCRT-II wHTH domain. These operons may be further extended with additional copies of Snf7 genes and other genes coding for a TM protein and an ABC ATPase. We also observed that a related VPS4-like AAA$^+$-ATPase was transferred from Archaea to Bacteria and is found in Cyanobacteria, Bacteroidetes, Verrucomicrobia, Nitrospirae, and Planctomycetes (*e.g.,* ACB74714.1, Opitutus; **Table S3**). In these operons, the *snf7* gene is displaced by an unrelated gene coding for a larger protein with TPR repeats followed by a 6TM domain, again suggesting a membrane-proximal complex.

Our analysis also showed that the bacterial PspA (*e.g.,* AEY64321.1, Clostridium; **Table S3**) might occur with a distinct AAA$^+$-ATPase in various bacterial clades. The resulting protein (*e.g.,* AEY64320.1, Clostridium) has two AAA$^+$-ATPase domains (*e.g.,* CKH37208.1, *Mycolicibacterium*) in the same polypeptide, with the N-terminal version being inactive. This gene dyad also occurs with either a previously unidentified membrane-anchored protein with a divergent Snf7 domain (OGG56892.1; **Table S3**) and other coiled-coil or **α**-helical domain-containing proteins. Both PspA and the membrane-associated Snf7, along with the AAA$^+$-ATPase, may occur in longer operons with other genes coding for an ABC-ATPase, an ABC TM permease, and a solute-binding protein with PBPB and OmpA domains [*e.g.,* OGG56892.1; **Table S3**].

#### 2.3 PspA with PspM or Thioredoxin

The association of ClgR-HTH with PspAM is also confined to this RsmP family, suggesting that these are also determinants of the rod-shaped morphology of the cell. The PspN presence in the immediate operon of ClgR-HTH–PspAM (containing ClgR, PspA, PspM) is limited to a few mycobacteria (CCP45543.1, *M. tuberculosis* H37Rv), which have an N-terminal PspN_N (as defined below) and C-terminal DUF3046. The remaining ClgR-HTH–PspAM operons lack the fused PspN_N–DUF3046 protein and instead contain only the ancestral DUF3046 located three genes downstream (ABK71106.1, *Mycolicibacterium smegmatis*). The duplicated DUF3046 domain forms the intact ClgR-HTH–PspAMN operon only in the *M. tuberculosis* complex (6, 7). The presence of the same family of thioredoxin with a different family of PspA (typically, two copies) in Cyanobacteria suggests that the thioredoxin homolog is involved in a similar redox activity to control PspA [AFZ14666.1, Crinalium; **Fig. 3**; **Table S3**].

#### 2.4 Novel contexts containing Toastrack

##### Toastrack and TM domains

In most homologs, we find that Toastrack domains are fused to N-terminal single or multi-TM domains such as PspC, LiaI-LiaF-TM, HAAS, SHOCT, strongly suggesting that the Toastrack domains are predominantly intracellular with N-terminal membrane tethers [**Fig. 4, 5**; **Table S4**]. In Cyanobacteria, we find variable multidomain proteins with an N-terminal TM anchor followed by a region containing the Toastrack domain flanked by immunoglobulin (Ig) and one or more catalytic domains such as a fringe-like glycosyltransferase or a caspase-like thiol peptidase [**Fig. 4**; **Table S4**; AFY83227.1, *Oscillatoria*; **Table S2**]. Further, in several architectures, the N-terminal TM regions fused to the Toastrack domain are replaced by at least two variants of the bihelical SHOCT (*e.g., Bacillus subtilis yvlB*, CAB15517.1) [**Fig. 4, 5**; **Tables S1** and **S4**]. We call these variants **SHOCT-like** domains to distinguish them from the classical SHOCT domain, as these include a domain partly detected by the Pfam **DUF1707** (8) model and another that has not been detected by any published profile. The SHOCT and related domains are fused to disparate domains and are typically found at the N- or C-termini of proteins.

##### Toastrack and transcription factors

We also discovered several conserved genomic contexts containing Toastrack, with likely roles in membrane-linked stress response: The first of these found across diverse bacterial lineages contains a core of four genes coding for i) a sigma factor, ii) a receptor-like single TM protein with an intracellular anti-sigma-factor zinc finger (zf-HC2, PF13490 in Pfam) and extracellular HEAT repeats, iii) one or two membrane-anchored Toastrack-containing proteins (AFK03672.1 Emticicia; **Table S4**), and iv) a previously uncharacterized protein with hits to the Pfam model DUF2089. We found that this Pfam model **DUF2089** can be divided into an N-terminal ZnR, central HTH, and C-terminal SHOCT-like domains (ADE70705.1, *Bacillus*) [**Fig. 4**; **Tables S2** and **S4**]. In a few of these operons, the membrane anchor of the Toastrack domain is a LiaI-LiaF-TM domain [**Fig. 5**; **Table S4**]. Variants of this system include additional genes coding for a protein with a LiaI-LiaF-TM domain fused to an N-terminal B-box domain (*e.g.,* ACO33311.1, *Acidobacterium*) or a PspC protein (*e.g.,* OGF50123.1, *Candidatus Firestonebacteria*) [**Fig. 5**; **Table S4**]. We propose that this three-gene system functions similarly to the classical *lia* operon in transducing membrane-associated signals to a transcriptional output affecting a wide range of genes via the sigma factor.

Similarly, an operon observed predominantly in various Proteobacteria and Bacteroidetes couples a protein with a membrane-anchored Toastrack domain (typified by AAM36414.1, *Xanthomonas*) with genes coding for an ABC-ATPase, a permease subunit, and a **GNTR-HTH** transcription factor with distinct C-terminal α-helical domain and another [**Fig. 5**; **Tables S2** and **S4**]. These operons also code for a previously uncharacterized protein matching the Pfam DUF2884 model. We show that these proteins are membrane-associated lipoproteins (*e.g.,* AJI31452.1, *Bacillus*), which might function as an extracellular solute-binding partner for the ABC-ATPase and permease components. A comparable operon found in Actinobacteria replaces the **GNTR-HTH** transcription factor with a ribbon-helix-helix (**RHH**) domain protein. In some Actinobacteria, the Toastrack domain encoded by the operon is fused to a SHOCT-like domain and is encoded adjacent to genes specifying a two-component system (CCP43715.1, *Mycobacterium*) or a transport operon (CAB88834.1, *Streptomyces*) [**Fig. 5**; **Table S4**]. These operons with the Toastrack domains are likely to couple transcriptional regulation to the sensing of membrane-proximal signal and transport [**Fig. 5**; **Table S4**]. The GNTR-HTH and RHH operons in these systems are likely to function as transcriptional regulators analogous to PspF and ClgR transcription factors from classical PSP systems.
